# Total Biosynthesis of *Pseudomonas aeruginosa*-Derived Azabicyclocarbamates Identifies Distinct Dehydrating Condensation Family Proteins

**DOI:** 10.64898/2026.07.17.738954

**Authors:** Xiaohan Liu, Soumaya Najah, Andrea Calderari, Zhilai Hong, Sébastien Halary, Carine Lombard, Arnaud Bolard, Katy Jeannot, Arnaud Gruez, Kira J. Weissman, Yanyan Li

## Abstract

Bacterial azabicyclocarbamates and related pyrrolizidine alkaloids play important roles in microbial interactions, and are scaffolds of therapeutic potential. Their biosynthesis involves a bimodular non-ribosomal peptide synthetase (NRPS), as well as a Baeyer-Villiger monooxygenase and tailoring enzymes, the latter contributing to the structural diversification of these compounds. Azetidomonamide A, a core metabolite produced by the major human opportunistic pathogen *Pseudomonas aeruginosa*, is a rare 4,7-bicyclocarbamate involved in modulating bacterial virulence that belongs to a unique family of natural products targeting ClpP proteases. In this study, we elucidated the full set of reactions leading to the 7-membered cyclocarbamate warhead, and reconstituted *in vitro* the biosynthesis of azetidomonamide A. Notably, this approach allowed for detailed characterization of a condensation (C) domain-catalyzed online dehydration via chemical capture of NRPS-tethered intermediates. Furthermore, we identified the dehydratase AzeD as the founding member of a distinct group of standalone proteins of the C domain family. Via combined structural, docking and biochemical analyses, we provided evidence that AzeD’s catalytic mechanism is distinct from that of dehydrating C domains, further expanding the known chemistry of these key biosynthetic enzymes.

## INTRODUCTION

Microorganisms produce a wide range of specialized metabolites, whose structures are tailored for their roles in the native producer within a specific environmental context,^1^ notably including host-pathogen interactions. The recently-discovered azetidine-containing alkaloids, the azetidomonamides,^2^ are core metabolites of the major human opportunistic pathogen *Pseudomonas aeruginosa* that have multifaceted impacts on the pathogen’s biology. Azetidomonamide A **1** is a rare azabicyclo[5,2,0]carbamate, whereas azetidomonamide B **2** (also termed azabicyclene)^3^ features an azetidopyrroline scaffold (**Figure 1**). The *aze* pathway is an integral part of a quorum-sensing regulated network,^4^ and is implicated in both virulence-related behaviors and host colonization.^2,5^ Recently, **1** but not **2**, was demonstrated to promote biofilm formation and pigment production in *P. aeruginosa*.^6^ Moreover, the 7-membered cyclocarbamate motif found in **1** was shown to act as a warhead, specifically targeting bacterial ClpP proteases by covalent modification of the active sites.^7^ The ClpP protease is a validated antibiotic target, as it is involved in pathogen protein homeostasis, stress response and virulence.^8,9^ These findings together suggest that **1** accounts for the virulence modulating effect of the *aze* pathway, and identify its scaffold as a promising lead for new antibacterials acting as ClpP inhibitors.

**Figure 1.**
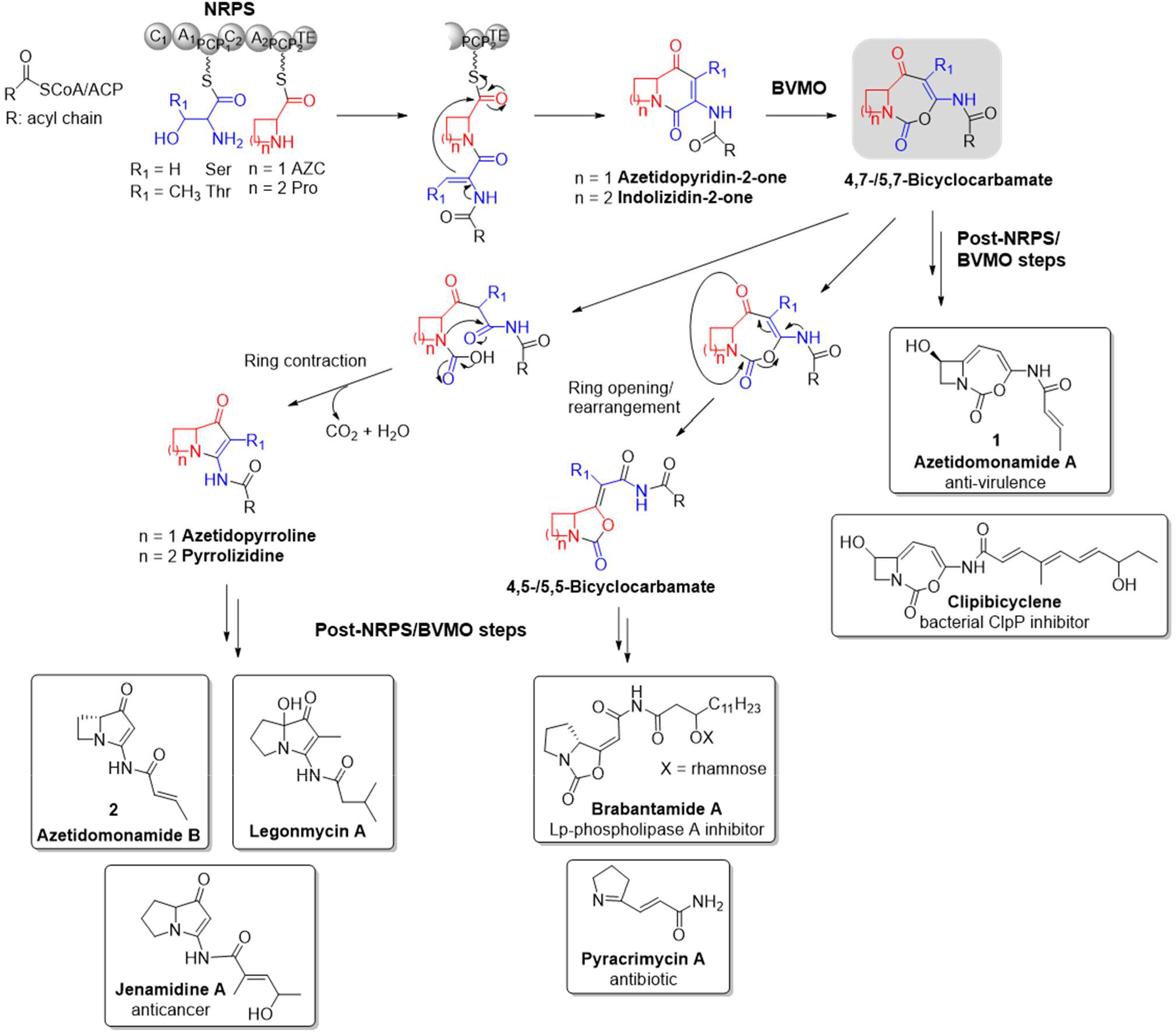
Generalized scheme for the biosynthesis of bacterial azabicyclocarbamates and related pyrrolizidine alkaloids. Structures of representative compounds are shown and their bioactivities, if known, are indicated. The common 7-membered bicyclocarbamate intermediate at the branching point is highlighted in grey. Key to abbreviations: NRPS, nonribosomal peptide synthetase; BVMO, Baeyer-Villiger monooxygenase; ACP, acyl carrier protein; C, condensation; A, adenylation; PCP, peptidyl carrier protein; TE, thioesterase.

The biosynthesis of azetidomonamides, like that of bacterial pyrrolizidine alkaloids and related bicyclocarbamates such as the brabantamides, invariably involves a bimodular non-ribosomal peptide synthetase (NRPS) and a Baeyer-Villiger monooxygenase (BVMO)^2,10–15^ (**Figure 1**). NRPSs are gigantic multienzymes composed of modules which each harbors a series of functional domains. NRPS modules minimally include adenylation (A) domains for selection and activation of amino acid substrates, condensation (C) domains that catalyze peptide bond formation, and peptidyl carrier protein (PCP) domains that serve as covalent supports for the substrates/chain extension intermediates. In the case of pyrrolizidine alkaloids and azetidomonamides, the NRPS concatenates an acylated Ser or Thr with a Pro or azetidine 2-carboxylic acid (AZC), followed by an online dehydration step to release the indolizidinone or azetidopyridinone product **3** (**Figures 1** and **2**). The BVMO then inserts oxygen to form a 7-membered cyclocarbamate. The fate of this common intermediate then diverges, as it can undergo ring opening coupled to decarboxylation, leading to ring contraction or rearrangement. Alternatively, additional modifications by a set of accessory enzymes yield **1** and the closely-related clipibicyclene (**Figure 1**), via a series of tailoring steps whose enzymology has remained largely elusive.^7^

Here, we leveraged total biosynthesis *in vitro* with the azetidomonamide pathway enzymes AzeB—G to elucidate the full route to azetidomonamide A **1**, and used the same system to prepare its Pro analog **1′**. The critical online dehydration of the Ser-bearing chain extension intermediate by AzeB was investigated in detail via chemical off-loading. Moreover, using an integrative structural biology approach, we identified AzeD as a novel type of stand-alone condensation domain catalyzing dehydration, and provided evidence for its catalytic mechanism. Overall, the reported results lay the foundation for engineering this unique family of therapeutically promising ClpP-targeting natural products^7^ that play key roles in bacterial biology.

## RESULTS AND DISCUSSION

### Functional Assignment of Enzymes Involved in Ring Modification

In addition to the NRPS (AzeB) and BVMO (AzeC), the *aze* biosynthetic gene cluster in *P. aeruginosa* encodes a standalone C domain-containing protein (AzeD), a short chain dehydrogenase (AzeE), a cytochrome P450 (AzeF), and a homologue of PhzA/B (AzeG) of the phenazine pathway belonging to the nuclear transport factor 2-like superfamily,^16^ as well as other proteins involved in the transport and the synthesis of NRPS precursors (**Figure 2A**).^2,3^ Based on this complement of enzymes, we reasoned that starting from the intermediate 4,7-bicyclocarbamate **4** (**Figures 1** and **2C**), AzeE would reduce the keto group to a hydroxyl to yield **5**, followed by dehydration to afford azetidomonamide C **6**.^6^ Both AzeD and AzeG were potential dehydratase candidates, as some C domains are known to have non-canonical dehydrating activities.^17,18^ Regio- and stereospecific hydroxylation of **6** by AzeF would then lead to **1**, the only reaction which had been characterized *in vitro* prior to this work.^6^

**Figure 2.**
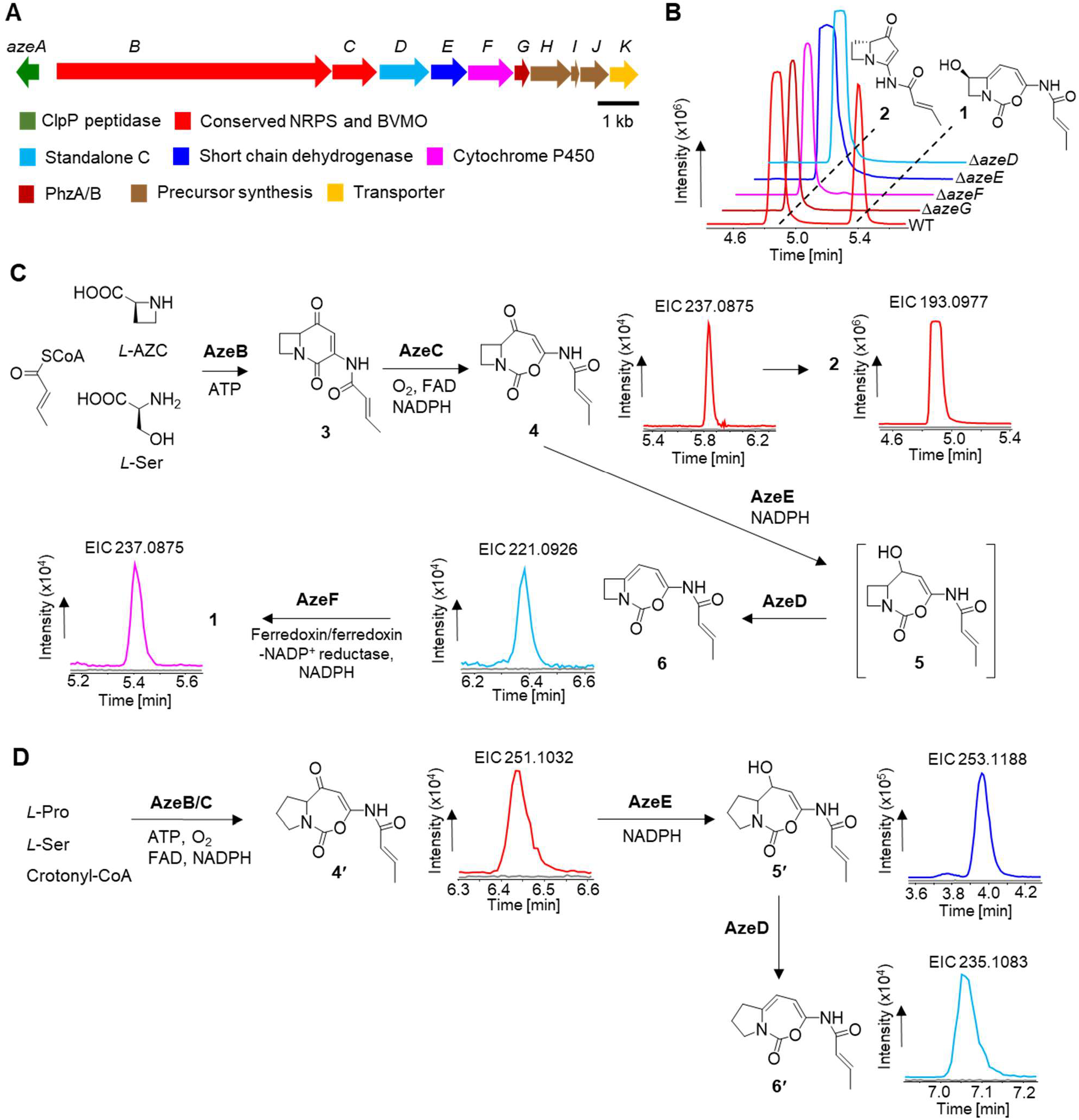
Characterization of ring modification reactions leading to azetidomonamides. **A**) Gene organization of the *aze* biosynthetic gene cluster in *P. aeruginosa*. **B**) Production of azetidomonamide A **1** and B **2** by the wild-type (WT) and single gene-deletion mutants. Combined extracted ion chromatograms (EICs) of [M+H]^+^ ions 237.0875 and 193.0977 corresponding to **1** and **2**, respectively, are shown. **C**) Step-wise reconstitution *in vitro* of azetidomonamide biosynthesis. The parentheses around **5** indicate that it was not directly detected. **D**) Characterization *in vitro* of AzeE and AzeD using Pro as substrate, leading to the detection of the key intermediate **5′**. For panels **C** and **D**, EICs corresponding to the expected product [M+H]^+^ ions are shown next to the compound structure. Colored curve, reaction; grey curve, heat-inactivated enzyme control. Accompanying UV and MS/MS spectra of the detected compounds are provided in **Figures S7**—**S14**).

To assign functions to these genes, single deletion mutants (Δ*azeD*, Δ*azeE*, Δ*azeF* and Δ*azeG*) of *P. aeruginosa* AB16.2 were generated (**Table S1**).^2^ In order to detect intermediates produced at only low titers, we initially optimized the culture conditions. Consistent with their roles in stimulating biofilm formation, the production levels of the azetidomonamides, and in particular that of **1**, were observed to increase significantly in static cultures in a previously-identified ‘optimal’ medium (2×M9) (**Figure S1**).^6^ Crude extracts of the strains grown under the biofilm condition were then analyzed by both UV absorbance and liquid chromatography coupled to high resolution mass spectrometry analysis (LC-HRMS) (**Table S3**), as characterization of the detected intermediates by NMR was hampered by their low yields.

In all four mutants, the production of **1** was abolished as expected, whereas **2** was synthesized at a comparable yield to that of the parental strain (**Figure 2B**). The (4,7)-bicyclocarbamate **4** was detected in Δ*azeD*, Δ*azeE* and Δ*azeF* mutants, albeit in low amounts (**Table S4** and **Figure S2**). Furthermore, azetidomonamide C **6** accumulated significantly in the Δ*azeF* strain, consistent with the established function of AzeF (**Figure S3**), but was not otherwise present systematically (**Table S4**).^6^ By contrast, **4** and **6** were barely detectable in the Δ*azeG* mutant. We also observed the equivalent Pro-containing intermediates in the mutant strains (**Figures S4**—**S5**), consistent with our previous finding that the *aze* pathway also naturally produces Pro-incorporated compounds **1′** and **2′** (the ′ indicates derivatives in which Pro replaces AZC).^2^ Cumulatively, these results showed that AzeD—G are all required for the full series of post-BVMO modifications *in vivo*.

To probe directly the function of each enzyme, recombinant proteins AzeB—G were produced individually in *Escherichia coli* and purified to homogeneity for characterization *in vitro* (**Figure S6**). Reactions were performed in a step-wise manner and their products were monitored by LC-HRMS, with product assignments supported by comparison to the metabolites observed *in vivo* (**Table S3**). Incubation of AzeB and AzeC in the presence of L-AZC and L-Ser with necessary cofactors, yielded **2** as a major product and a small amount of **4** (**Figures 2C** and **S7**). Time course analysis showed that **2** was already dominant after 1 min (**Figure S8**), demonstrating the intrinsic reactivity of the 7-membered cyclocarbamate. Next, AzeE was added in the presence of excess NADPH, but no hydroxyl product **5** was detected, even with variation of the assay conditions (e.g. pH, incubation time, enzyme concentration, etc.).

We reasoned that the failure to observe **5** derived from the intrinsic instability of the strained azetidine ring, as **5** was not observed *in vivo*. To test this idea, *in situ* enzymatically generated **4′** from Pro was used as substrate for AzeE (**Figures 2D** and **S9**). Gratifyingly, a new product with [M+H]^+^ 253.1184 (r.t. 4 min, *m/z* calc’d for C_12_H_17_N_2_O_4_ = 253.1188, Δ ppm =−1.6) corresponding to **5′** was detected in the reaction, but not in the control (**Figures 2D** and **S10**). Subsequent addition of AzeD yielded a compound with [M+H]^+^ 235.1082 (r.t. = 7.05 min, *m/z* calc’d for C_12_H_15_N_2_O_3_ = 235.1083, Δ ppm = −0.4) consistent with **6′**, as identified in the Δ*azeF* strain (**Figures 2D** and **S11**). In contrast, incubation with AzeG did not lead to the dehydrated product **6′**. Thus, by working with the Pro series of compounds instead of the AZC derivatives, we were able to conclusively identify AzeE as a ketoreductase and AzeD as a dehydratase. The stereochemistry of the ketoreduction remains to be determined, as it is masked by the subsequent dehydration.

Next, the cytochrome P450 AzeF was assayed with the *in situ*-generated **6′** and led to the production of **1′**, as expected (**Figure S12**). Reasoning that the downstream enzymes might overcome the stability issues with **5**, we then performed a one-pot reaction with all five enzymes AzeB—F using AZC as substrate. This approach indeed resulted in synthesis of **1**, albeit at low yield, whose identity was confirmed by comparison to purified standard^2^ (**Figures 2C** and **S13**). The dehydrated product **6** was also observed in this case (**Figures 2C** and **S14**).

Taken together, these experiments demonstrated conclusively that the three enzymes AzeD—F are necessary and sufficient *in vitro* for full ring modification. The role of AzeG thus remains elusive, as although it is essential *in vivo*, its presence *in vitro* does not modify the product profile (**Figure S15**). This situation is reminiscent of PhzA/B from the phenazine pathway in *Pseudomonas*, which are also required only *in vivo*.^19,20^ A possible explanation for this discrepancy is that AzeG assists other pathway enzymes or suppresses side reactions, as proposed for other nuclear transport factor 2-like proteins,^16^ boosting the overall efficiency of the biosynthesis. This hypothesis could be evaluated via detailed kinetic study in future work.

**AzeB_C**_**2**_ **Catalyzes Online Dehydration**. The synthesis of the azetidopyridinone **3** by AzeB requires an online dehydration step of the acylated-Ser (**Figure 3A**). We hypothesized that this reaction was catalyzed by the AzeB_C_2_ domain that belongs to the “modified AA” (C_modAA_) subfamily.^21^ Identified members of this subfamily include AmbE_C_modAA_ (methoxyvinylglycine pathway) and AlbB_C_2_/C_3_ (albopeptide pathway) that act as dehydratases.^17,22,23^ To characterize the function of AzeB_C_2_ *in vitro*, full-length AzeB and several truncated forms were produced in *E. coli* as recombinant proteins (**Figures 3B** and **S6**). Chemical capture of NRPS-tethered intermediates was then performed using cysteamine as the off-loading agent.^24^

**Figure 3.**
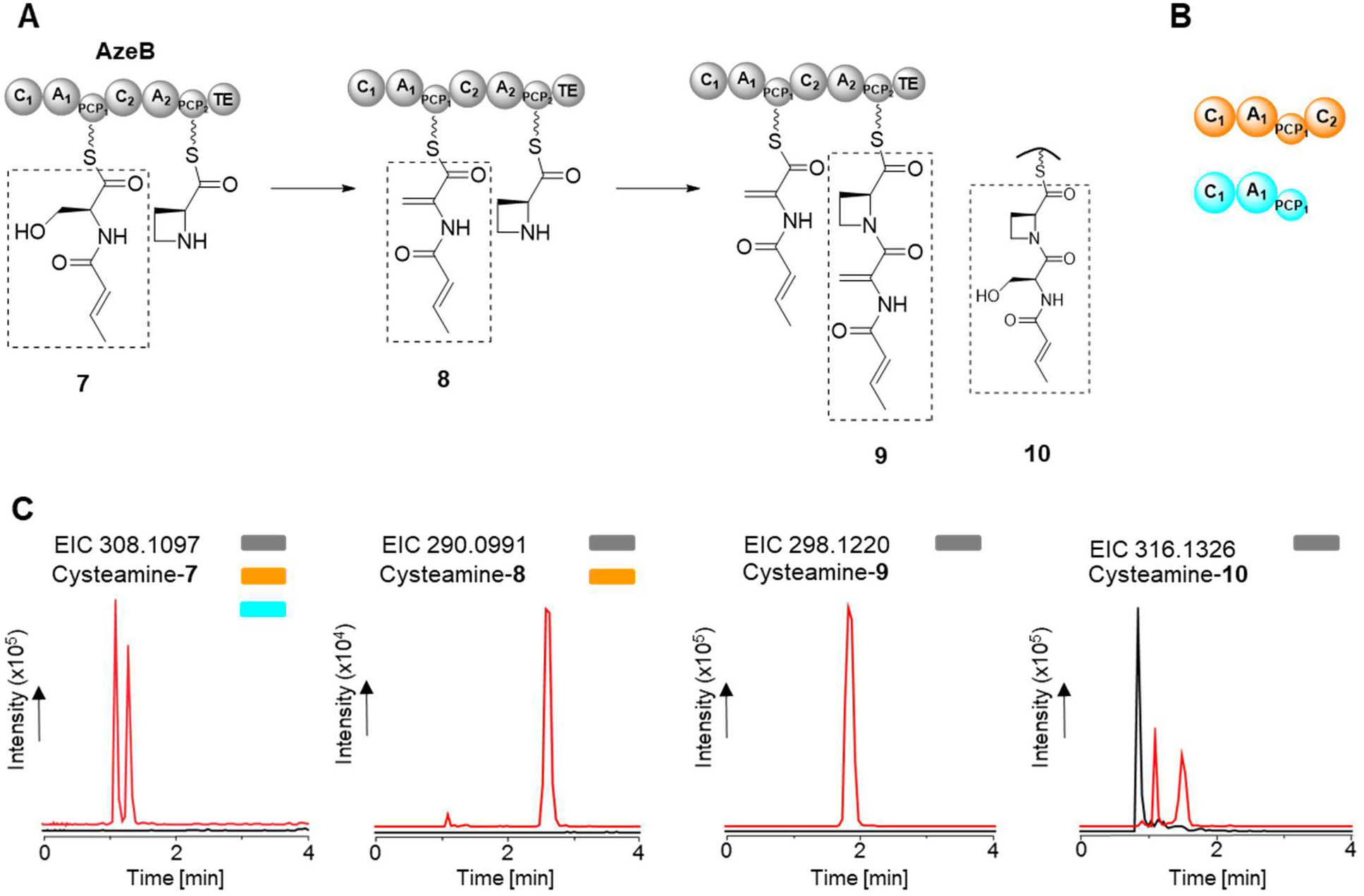
Characterization *in vitro* of the online dehydration step. **A**) Series of reactions catalyzed by the NRPS AzeB. **10** is a dead-end intermediate, as it cannot undergo chain release via cyclization. **B**) Schematic representation of the truncated versions of AzeB used in this study. **C**) Detection of AzeB-linked intermediates by cysteamine offloading.^24^ The EICs corresponding to the [M+H]^+^ ions are shown. Key to the reactions: red curve, active enzyme; black curve, heat-inactivated control. The colored bars indicate for which AzeB version the intermediate was detected. The structures, UV and MS/MS spectra of the cysteamine off-loaded compounds are provided in **Figures S16**—**S19**.

In the assay with full-length AzeB, crotonylated (Cr)-Ser **7**, Cr-dehydro-Ala (Dha) **8**, dipeptide Cr-Dha-AZC **9** and a very small amount of Cr-Ser-AZC **10** were detected (**Figures 3C** and **S16**—**S19**). When module 1 of AzeB was assayed (i.e., AzeB_C_1_-A_1_-PCP_1_), only Cr-Ser **7** was captured. By contrast, **8** was detected as a major product in the reaction with the tetradomain AzeB_C_1_-A_1_-PCP_1_-C_2_. Together, these experiments demonstrate that the AzeB_C_2_ domain dehydrates Cr-Ser tethered to PCP_1_, a timing in agreement with that established for dehydrating C_modAA_ domains.^17,23,25^

The detection of the dead-end intermediate **10** (**Figure 3C**) in the AzeB reaction is intriguing, as in principle, it could arise from promiscuous condensation activity by AzeB_C_2_ acting on Cr-Ser **7**, or given the intrinsic reactivity of Dha, rehydration of the dipeptide **9**.^26^ To discriminate between these possibilities, the AzeB reaction was conducted in H_2_^18^O. LC-HRMS analysis revealed that **10** was fully labeled with ^18^O (**Figure S19**), a result consistent with the re-hydration hypothesis. Globally, these data strongly argue that AzeB_C_2_ discriminates against Cr-Ser, instead only catalyzing ligation of the dehydrated form (Cr-Dha) with AZC (**Figure 3A**). This result contrasts with the relaxed condensation specificity reported for LynD_C_2_ involved in legonmycin biosynthesis, as LynD_C_2_ has been shown to accept both dehydrated and non-dehydrated substrates.^25^

### AzeD Belongs to A Distinctive Group of Standalone C Proteins

Having elucidated the full biosynthetic pathway to **1**, we were intrigued by AzeD, an unprecedented dehydratase belonging to the C domain family. Sequence analysis by the Natural Product Domain Seeker 2 (NaPDoS2) tool^27^ that is based on phylogenetic relationships to experimentally characterized C domains, did not reveal matches for AzeD, underscoring its novelty. Moreover, in AzeD, the first His in the HHxxxDG motif of canonical C domains^28^ is replaced by Pro. To gain insight into the prevalence and distribution of AzeD, we carried out a protein BLAST against the UniREF_90 database followed by an HMMsearch using stringent parameters, revealing 35 AzeD homologs (**Figure S20**). These sequences, along with AzeB_C_2_ and a validated C domains dataset used previously for an evolution study,^29^ were employed for phylogenetic analyses. The resulting unrooted tree exhibits a structure similar to that observed previously (**Figure 4**),^29^ with deep branches supported by high bootstrap values (Transferred Bootstrap Expectation > 75%). AzeB_C_2_ is positioned in the C_modAA_ clade, as expected, whereas AzeD and its homologues group into a single clade, close to those of heterocyclizing and epimerizing enzymes. As all AzeD homologues are encoded in the genomes as standalone proteins, we have named this newly-identified clade C_standalone_. Approximately 1/3 of the C_standalone_ members display variations in the first His of the conserved catalytic motif, while 1/3 are encoded within a genomic neighborhood containing NRPS and/or polyketide synthase genes, suggesting their involvement in natural product biosynthesis (**Figure S20**). It remains to be determined if additional or all C_standalone_ family members are dehydratases.

**Figure 4.**
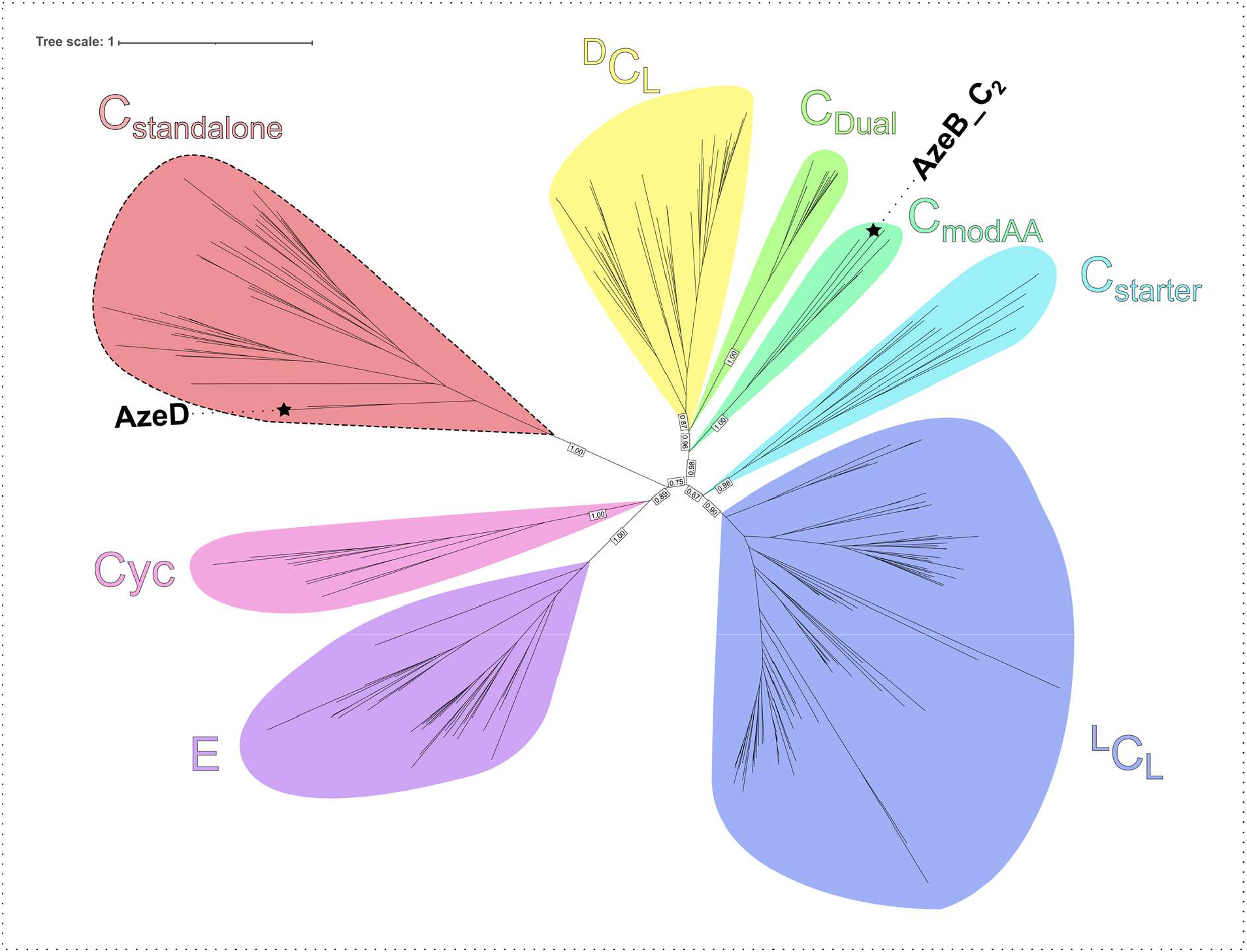
Unrooted phylogenetic tree of 246 C-domain sequences reconstructed using PhyML 3.0^48^ and Q.pfam substitution matrix with the decorations +R+F. Numbers correspond to branch support values >0.6 calculated from 100 replicates. Stars indicate AzeB_C_2_ and AzeD sequences. The clades are colored according to C-domain functions: Epimerization (E), Heterocyclization (Cyc), Starter (C_Starter_), Dehydroamino acid associated (C_modAA_), Dual epimerization/condensation (C_Dual_), condensation between L- and L-amino acids (^L^C_L_), and condensation between D- and L-amino acids (^D^C_L_). The newly identified salmon-colored clade corresponds to the C_standalone_ subfamily.

### Characterization of the AzeD Dehydratase by X-ray Crystallography and Small-angle X-ray Scattering

To deepen our understanding of AzeD, we aimed to solve its crystal structure at high resolution. Recombinant AzeD was purified to homogeneity by two-step liquid chromatography (**Figure S21**). After confirming the monodispersity of the sample by DLS (**Figure S22**), we crystallized the *apo* form of AzeD by vapor diffusion and solved its structure at a resolution of 1.55 Å by molecular replacement using an AlphaFold2 model^30^ (PDB ID: 8S6L) (**Table S5**). The asymmetric unit contains a monomer of 48.1 kDa molecular weight, whose physiological relevance was confirmed by small-angle X-ray scattering (SAXS) (observed molecular weight: 43—50 kDa vs. calculated MW: 51.2 kDa)^31^ (**Figure S23** and **Table S6**). This result contrasts with the homodimeric structure previously described for the dehydrating AmbE_C_modAA_ domain (PDB ID: 7R9X),^22^ but is consistent with the monomeric character of NRPS multienzymes.^32^ Certain residues are missing in the electron density maps, including the N-terminal His_6_ tag and the adjacent Met1, as well as a series of amino acids (Arg435—Gln441) located in the C-terminal loop, attesting to their disordered character. Overall, 98.1% (423) of the residues are situated in the favored region of the Ramachandran plot, and 98.0% (344) display favored rotamers. Several side chains exhibit multiple conformations, and two of the 33 prolines (Pro108 and Pro277) adopt a *cis* conformation.

As with other C domains, AzeD likely evolved from an ancestral chloramphenicol acetyltransferase (CAT) through gene duplication, fusion, and maturation of a protomer into an overall pseudo-dimer. In the solved structure, the N- and C-terminal lobes are separated by a central opening, giving rise to an overall V-shaped arrangement (**Figure 5A**).^33,34^ The N-terminal sub-domain comprises a central six-stranded β-sheet (β4-β5-β6-β1-β11-β12), two of which (β11-β12) are contiguous and form a β-crossover, along with four α-helices and two peripheral antiparallel β-strands (β2-β3). The C-terminal sub-domain is composed of a β-sheet containing six twisted β-strands(β7-β14-β13-β10-β8-β9) flanked by six α-helices. The lobes are connected via conserved structural elements: helix α5, a 3_10_-helix situated in the N-terminal region (α-crossover), and a loop-β-strands-loop motif (β-crossover) (29 residues, Ile359—Tyr387) located between the β10 and β13 strands^35^ (**Figure 5A**).

**Figure 5.**
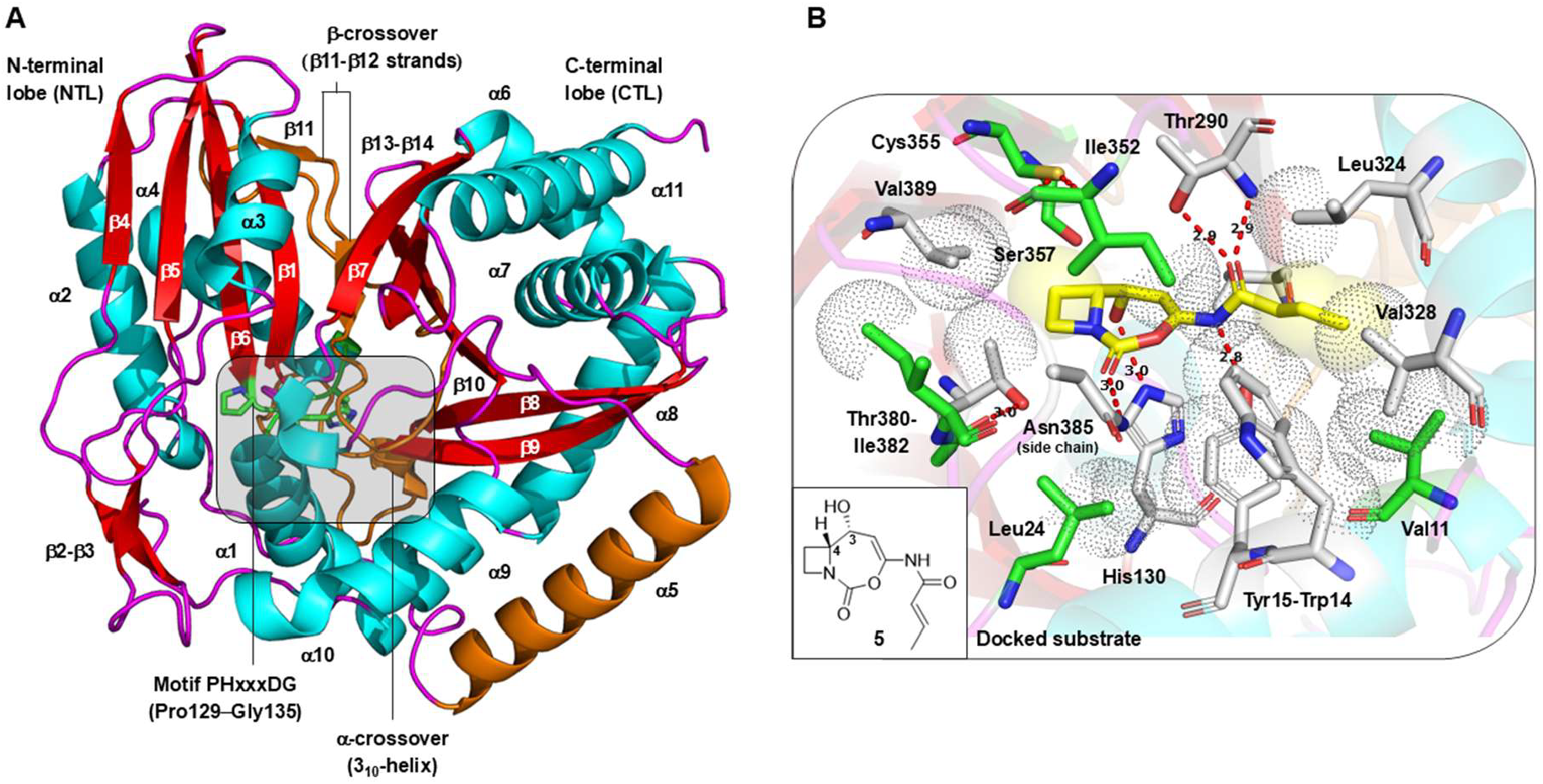
Crystal structure of the AzeD dehydratase solved at 1.55 Å resolution (PDB ID: 8S6L) and computational model for substrate binding. **A**) The AzeD monomer consists of N- and C-terminal lobes separated by a central opening leading to an overall V-shaped architecture. The α-helices and β-strands are represented in cyan and red, respectively. The interlobe structural elements (helix α5 and α-/β-crossovers) are depicted in orange, and the catalytic motif is shown in green. **B**) Binding mode of the substrate **5** (yellow) predicted by molecular docking using the CB-Dock2 program,^49^ depicting the stereoisomer of **5** with the *R* configuration at C-3. This analysis revealed a series of hydrophobic interactions (spheres) and hydrogen bonds (dotted lines) to the azetidopyroxydinol heterocycle and butenamide moiety, which position the ligand appropriately relative to the catalytic His130. Residues directly involved in recognition of the docked substrate (i.e., Trp14, Tyr15, His130, Val288, Thr290, Leu324, Val328, Thr380, Asn385 and Val389) are indicated in gray, while those engaged in interactions with other residues (i.e., Val11, Leu24, Ile352, Cys355 and Ile382) are displayed in green.

To further probe the AzeD structure, we compared it to that of the closest homologues identified using the Dali server, and in particular, NRPS-integrated C domains with classical condensation (Txo2_C1, PDB ID: 6P1J)^35^ or alternative dehydration functions (AmbE_C_modAA_, PDB ID: 7R9X)^22^ (**Figure S24**). These domains all adopt an overall CAT fold, but AzeD more closely resembles the canonical C domain (RMSD to Txo2_C1 = 4.1 Å (243 atoms)) than the variant capable of amino acid dehydration (RMSD to AmbE_C_modAA_ = 6.3 Å (299 atoms)). Furthermore, AzeD differs from both homologues in terms of the amplitude of interlobe opening, and the position, orientation, number and relative length of several secondary structures, especially bridging elements (**Figures S25** and **S26**).

### Active Site Organization and Proposed Catalytic Mechanism

The second His in the conserved HHxxxDG motif in canonical C domains^28^ has been previously proposed to either act as a catalytic general base by deprotonating the α-amine of the acceptor amino acid during peptide bond formation,^36,37^ or to participate in substrate positioning.^38^ However, more recent work demonstrated its role in stabilizing the condensation transition state by acting as a hydrogen bond acceptor.^39^ In AzeD, this catalytic His is conserved (His130), but the directly upstream His is replaced by a Pro (P^129^HxxxDG^135^) (**Figure S26**). Concerning substrate **5**,neither the stereochemistry at C3 or C4 is known. Thus, to obtain mechanistic insight into the dehydration reaction and in particular the potential role of His130, we carried out *in silico* docking experiments between AzeD and the four possible stereoisomers of **5**, using the CB-Dock2 program.

This analysis revealed that the active site is situated at the interlobe interface (**Figures 5** and **S27**). Interestingly, the presence of Pro129 introduces a bend in the conserved active site motif that places His130 within hydrogen bonding distance of the leaving C3-OH group (3.0 Å) in the docked model, but only when C3 exhibits *R* stereochemistry (**Figures 5B** and **S28**). According to these calculated poses, substrate recognition is mediated by the side chains of certain residues, notably Trp14, Leu324, Val328, Thr380 and Val389, which participate in hydrophobic interactions with the ligand, while Tyr15, Thr290, and Asn385 make hydrogen bonds with the amine or carbonyl groups. However, when the ligand has the opposite *S* stereochemistry at C3, the resulting orientation within the active site no longer permits a stabilizing interaction between His130 and any polar functional group of **5** (**Figure S28**). Taken together, these data suggest that stereospecific reduction by AzeE results in the *R* configuration at C3.

Amino acid dehydration catalyzed by C_modAA_ domains likely occurs via E1cB-type chemistry. In the proposed mechanism, α-proton abstraction catalyzed by the catalytic His results in an enolate intermediate, for which stabilization of the negative charge is afforded by the adjacent thioester^22,29,40^ (**Figure S29A**). In the case of the AzeD substrate **5**, no comparable stabilization is available to the anion. In this context, we propose that the reaction proceeds in E1 fashion (**Figure S29B**). Specifically, His130 would first act as a general acid to protonate the C3-OH, a function potentially enabled by the adjacent His-to-Pro substitution, with the departure of water facilitated by the neighboring conjugated system. His130 would then serve as a catalytic general base to remove the C4 proton, leading to the final, extended conjugated system. Alternatively, formation of the double bond and removal of water could take place in concerted fashion (E2 mechanism) (**Figure S29C**). This reaction would occur with the proton (C4-H) and hydroxyl (C3-OH) arranged *anti*-periplanar to each other, and thus necessitate the *R* stereochemistry at the C4 center that is consistent with our docking analysis.^41^

Due to its proximity, we reasoned that Cys355 could function alongside His130 to affect this chemistry, but Thr290 and Ser357 were also considered as potential catalytic residues. Thus, to discriminate between the E1 and E2 mechanisms, we generated alanine mutants of AzeD residues His130, Thr290, Cys355 and Ser357. Analysis of the resulting purified proteins (**Figure S30**) by circular dichroism (CD) revealed that all of the mutations introduced structural perturbation, but to differing extents, with the H130A most impacted (**Figure S31**). Indeed, the H130A was completely inactive (**Figure S32**), which is consistent with its essential catalytic role, but which may also be partly attributable to unfolding. However, the three other mutants retained essentially wild-type dehydration activity (**Figure S31**), despite the introduced structural changes. As only mutation of His130 impacted AzeD activity, these data favor the proposed one-base E1 mechanism.

## CONCLUSIONS

In this work, we reconstituted *in vitro* the full biosynthetic pathway to 7-membered bicyclocarbamate azetidomonamide A **1** and its Pro analogue **1′**. Given that these enzymes operate on a substrate located at a key branch point (**Figure 1**), introducing them into related pyrrolizidine alkaloid pathways,^10–15^ could provide access to novel ClpP-targeting molecules. This work thus lays the foundation for genetically engineering these complex scaffolds of both ecological^2,6,7^ and therapeutic importance.^10,42–46^

From a fundamental perspective, it is remarkable that the biosynthesis of **1** involves both an NRPS-embedded and a standalone C domain for dehydration. While AzeB_C_2_ fits into the previously identified C_modAA_ phylogenetic group,^21^ AzeD belongs to the newly discerned, noncanonical C subfamily, whose members are all discrete proteins (C_standalone_). Structural, docking and biochemical analyses of AzeD are consistent with *R* stereochemistry at the C3-OH center of its substrate **5**, and E1-type dehydration. As this mechanism differs from that proposed for C_modAA_ domains functioning on NRPS-tethered peptides,^22,29^ these findings expand the functional scope of non-canonical C domains/proteins, highlighting them as a driving force for diversification of NRPS products.^33,47^ The convergent evolutionary history of AzeD and AzeB_C_2_-type C domains towards a shared dehydration function warrants further investigation.

## ASSOCIATED CONTENT

Supporting Information

Detailed description of materials and methods used and supplementary figures and tables (PDF).

## ACKNOWELDGEMENTS

This work was funded by the Agence National de la Recherche (ANR) to Y. L and K. J.W. (ANR20-CE92-0038-02 NRPSBacAza). X.L was supported by the China Scholarship Council (CSC), within the framework of the CSC-Sorbonne University program. Dr. A. Marie and R. Puppo in the MNHN Bioorganic analytical platform are thanked for their assistance with the mass spectrometry analysis. We are also grateful to Dr. M. Savko and Dr. W. Shepard for help with diffraction data acquisition on the beamline PROXIMA 2A (SOLEIL synchrotron).

