## Supplementary material for "Total Biosynthesis of *Pseudomonas aeruginosa*-Derived Azabicyclocarbamates Identifies Distinct Dehydrating Condensation Family Proteins": Materials and Methods; Suplementary Tables and Figures

[a]Unit Molecules of Communication and Adaptation of Microorganisms (MCAM), UMR 7245 CNRS-Muséum National d'Histoire Naturelle (MNHN), 75005 Paris, France

[b]Université de Lorraine, CNRS, IMoPA, F-54000 Nancy, France

[c]Laboratoire de Bactériologie, Centre National de Référence (CNR) de la Résistance aux Antibiotiques, Centre Hospitalier Régional Universitaire (CHRU) de Besançon, UMR6249 'Chrono-Environnement', Boulevard Fleming, 25030 Besançon, France

[§]Current address: Génomique Métabolique, Genoscope, Institut François Jacob, CEA, CNRS, a Université Evry, Université Paris-Saclay, 91057 Evry France.

[%]Current address: Smaltis, 25000 Besançon, France.

[£]Current address: CAS Key Laboratory of Quantitative Engineering Biology, Shenzhen Institute of Synthetic Biology, Shenzhen Institute of Advanced Technology, Chinese Academy of Sciences, Shenzhen 518055, China.

[#]These authors contributed equally

*Correspondence should be addressed to:


### Materials and Methods

#### Bacterial strains, plasmids and growth conditions

All chemicals and biochemicals were purchased from Merck or Carl Roth, while molecular biology reagents were purchased from New England Biolabs, unless otherwise stated. Bacterial strains and plasmids used in this study are listed in **Table S1**. *Escherichia coli* and *Pseudomonas aeruginosa* strains were routinely maintained in Lysogeny Broth (LB)-Miller medium at 37 °C. For gene deletion in *P. aeruginosa*, strains were grown in Mueller Hinton Broth (MHB) with adjusted concentrations of divalent cations $Ca^{2+}$ and $Mg^{2+}$ (Beckton Dickinson), or on Mueller Hinton Agar (MHA) (Bio-Rad). For metabolic profiling, *P. aeruginosa* and mutants were grown in 2×M9 medium,[1] which contains (per liter): 7.0 g $Na_2HPO_4$, 3.5 g $KH_2PO_4$, 1 g NaCl, 1 g $NH_4Cl$ and 10 mL of glucose (1% w/v), supplemented with 2 mL of 1 M $MgSO_4$, 0.6 mL of 1 M $CaCl_2$ and 10 mL trace elements solution (100× trace elements solution (per liter): 5 g EDTA, 0.83 g $FeCl_3 \cdot 6H_2O$, 84 mg $ZnCl_2$, 13 mg $CuCl_2 \cdot 2H_2O$, 10 mg $CoCl_2 \cdot 2H_2O$, 10 mg $H_3BO_3$, 1.6 mg $MnCl_2 \cdot 4H_2O$). For protein expression, certain *E. coli* strains were grown in Terrific Broth (TB) medium (per liter: 12 g tryptone (Gibco), 24 g yeast extract (Gibco), 4 mL glycerol, 100 mL solution containing 2.31 g $KH_2PO_4$ and 12.54 g $K_2HPO_4$).

#### Generation of *P. aeruginosa* mutants

During the course of investigating the function of AzeA in azetidomonamide biosynthesis,[2] gene *azeA* (PA3326) was inactivated to generate *P. aeruginosa* AB16.2ΔazeA. Metabolic profiling by LC-HRMS did not reveal any difference in azetidomonamide production between *P. aeruginosa* AB16.2ΔazeA and the AB16.2 strain. Single gene deletion of *azeD* (PA3329), *azeE* (PA3330), *azeF* (PA3331) and *azeG* (PA3332) was then performed in *P. aeruginosa* AB16.2ΔazeA by homologous recombination, as previously described.[3,4] Briefly, two regions of 0.5 kb flanking the fragment to be deleted were amplified from genomic DNA, assembled by overlap PCR and cloned into the suicide vector, pKNG101.[5] Recombinant plasmids were introduced into *P. aeruginosa* by triparental conjugation using the mobilisation properties of the broad-host range helper plasmid pRK2013.[6] Transconjugants were selected on *Pseudomonas* Isolation Agar supplemented with streptomycin (2 000 mg/L). Excision of pKNG101 from transconjugants was achieved by selection on M9-sucrose medium. The chromosomal deletions were confirmed by PCR and DNA sequencing.

#### Construction of protein expression plasmids

Genes were amplified from the genomic DNA of the *P. aeruginosa* AB16.2 by PCR using Phusion High-Fidelity DNA polymerase and appropriate primers (**Table S2**). Standard restriction enzyme digestion and ligation procedures were used for cloning. For expression in *E. coli* of *azeB* (full length and truncated), *azeC*, *azeD*, *azeG* and *azeF*, the target gene was cloned between the *Nde*I and *Hind*III sites of pET28a(+). For *azeE* expression, the gene was cloned between the *Nde*I and *Xho*I sites of pET29b(+). For *azeD* expression for crystallization, the gene was cloned between the

*Bam*HI and *Hind*III sites of pET28a(+). All constructs were verified by sequencing (Eurofins).

**Protein expression and purification for *in vitro* assays**

*E. coli* BL21(DE3) was used for protein expression except in the case of AzeB and its truncated variants, which were produced in the *E. coli* BAP1 strain.[7] The latter encodes a copy of the broad-spectrum phosphopantheteinyl transferase (PPTase) Sfp from *Bacillus subtilis* in its genome,[8] and is suitable for producing assembly-line enzymes such as NRPSs in their *holo*-PCP forms. Recombinant *E. coli* strains harboring the expression plasmids were pre-grown overnight at 37 °C and subsequently inoculated into 1 L of LB medium (for expression of AzeB and truncated variants, AzeD and mutants, AzeE, AzeG and AzeF) or 2 L of TB medium (for expression of AzeC) supplemented with kanamycin (50 μg/mL). The cultures were grown at 37 °C with shaking until the optical density at 600 nm ($OD_{600}$) reached 0.7−1. The cultures were then cold shocked on ice for 30 minutes before induction with isopropyl-β-D-thiogalactopyranoside (IPTG) at a final concentration of 0.1 mM. Cells were allowed to grow at 16 °C for 20 h before being harvested.

All subsequent procedures were performed at 4 °C. The cell pellet was resuspended in 25 mL buffer A (50 mM Tris-HCl, pH 7.8, 300 mM NaCl, 10 mM imidazole, 10% glycerol) supplemented with 0.1 mg/mL DNAase I, followed by lysis by three passages through a French press, or by sonication (Sonifier™ SFX250, Branson). After centrifugation (48,000*g*, 45 mins), the supernatant was purified by nickel-affinity chromatography by gravity using 4 mL Ni-NTA agarose (50% slurry) (Qiagen) packed in a 12 mL fritted column (Merck Millipore). The resin was washed with water and pre-equilibrated with buffer A. After sample loading, the resin was washed using buffer A and proteins were eluted in a step-wise manner with increasing concentrations of imidazole (60, 100, 250 and 500 mM). After verification of purity by sodium dodecyl sulfate-polyacrylamide gel electrophoresis (SDS-PAGE), fractions containing the target protein were combined and concentrated using a Vivaspin 15R concentrator (Sartorius). The protein was then desalted using a PD-10 column (Cytiva), buffer exchanged into the storage solution (50 mM Tris-HCl, pH 7.8, 100 mM NaCl and 10% glycerol) and further concentrated. Aliquots of the purified protein were flash frozen in liquid $N_2$ and stored at −80 °C until use. From 1 L culture, about 5 mg AzeB, 9 mg AzeC, 8 mg AzeD or variants, 12 mg AzeE, 18 mg AzeF and 13 mg AzeG were obtained.

**Metabolic profiling of *P. aeruginosa* mutant strains**

All strains were inoculated and grown in 2 mL of LB medium in a 50 mL falcon tubes at 37 °C with shaking (180 rpm) for 16 hours. The pellets were harvested and washed with 10 mL 2×M9 medium followed by centrifugation (7,000*g*, 10 min). This wash procedure was repeated twice to completely change the medium. The pellets were then inoculated into 10 mL 2×M9 medium in 100 mL flasks, and incubated without shaking at 30 °C for 48 hours. This biofilm condition was shown to enable azetidomonamide **1** production at an improved yield. Each strain was measured in triplicate (biological replicates). The stationary culture was extracted with 3× 10 mL ethyl acetate. The organic layers were combined and evaporated to dryness. Crude extracts were resuspended in

80% acetonitrile and subjected to LC-HRMS analysis. It is worth noting that many compounds produced by the *aze* pathway have identical masses. Therefore, to assign their identities unequivocally, LC retention times and $MS^2$ fragmentation data were both systematically taken into account, and correlations made between compounds produced *in vivo* and in the enzymatic assays *in vitro*. When applicable, UV absorbance spectra were also compared to those reported in the literature.[1]

**LC-HRMS and MS/MS analysis**

LC-HRMS and $MS^2$ data were acquired on an ultra-high performance LC system (Ultimate 3000 RSLC, Thermo Scientific) coupled to a high-resolution electrospray ionization-quadrupole-time of flight (ESI-Q-TOF) mass spectrometer (MaXis II ETD, Bruker Daltonics). An Acclaim RSLC Polar Advantage II column (2.2 μm, 2.1 × 100 mm, Thermo Scientific) was used for LC separation at a flow rate of 0.3 mL/min. The gradient of solvent A (MilliQ water with 0.1% (v/v) formic acid) and solvent B (HPLC-MS grade acetonitrile with 0.1% formic acid) over a total run-time of 16 min was: 0−1 min: 5% B; linear increase from 5% to 100% B over 10 min; 11−12 min: 100% B; 12−13 min: linear decrease 100% to 5% B; 13−16 min: 5% B. Mass spectra were acquired in the *m/z* range 50−1300 in positive ion mode. The acquisition parameters of the ESI source were as follows: nebulizer gas 2.4 bar, dry heater 200 ºC, dry gas 8.0 L/min, capillary voltage 3500 V, end plate offset 500 V and charging voltage 2000 V. For MS/MS analysis, the auto mode (collision energy 32−48 eV) was chosen with the same parameters as the MS method. The data were treated with Data Analysis 4.4 (Bruker Daltonics).

**Cysteamine offloading of AzeB-tethered intermediates**

A standard reaction of AzeB or its truncated version was carried out with 1 mM crotonyl-CoA, 1 mM L-serine, 1 mM L-azetidine-2-carboxylic acid (AZC), 4 mM ATP and 24 μM enzymes in HEPES buffer (50 mM HEPES, 8 mM $MgCl_2$, pH 8.0) in a final volume of 50 μL. The reaction was incubated at 30 °C for 2 h, before addition of cysteamine-HCl (pH 7.0) solution to a final concentration of 50 mM, followed by incubation at 4 °C for 16 h. Next, the reaction was quenched by addition of 50 μL acetonitrile followed by freezing at −20 °C for 30 min to precipitate the proteins. The resulting supernatants were subjected to LC-HRMS analysis, as described earlier. For reactions in $H_2{}^{18}O$ (Eurisotop), enzyme stock solution and substrate mixture in HEPES buffer were freeze-dried and re-dissolved in the same original volume, and then combined. After incubation at 30 °C for 2 h, cysteamine-HCl in $H_2{}^{18}O$ solution was added and offloading assays followed the same procedure as described above.

**Enzymatic assays *in vitro***

<u>With Pro as substrate</u>: The reaction was conducted with 1 mM crotonyl-CoA, 1 mM L-serine, 1 mM L-proline, 4 mM ATP and 10 μM AzeB in buffer (50 mM HEPES, 8 mM $MgCl_2$, pH 8.0) in a final volume of 100 μL and incubated at 30 °C for 2 h. Twelve μL of this reaction were removed,

quenched with an equal volume of acetonitrile and stored for later LC-HRMS analysis to detect AzeB products. In parallel, a negative control was prepared containing boiled AzeB in the same enzyme/substrate ratio, in a final volume of 25 μL. Next, 4 mM NADPH and 2 mM FAD were added to the remaining 88 μL of AzeB reaction to prepare the AzeC pre-reaction, in a final volume of 100 μL. Twelve μL of AzeC pre-reaction were then removed and incubated with boiled AzeC as the negative control. AzeC (15 μM) was added to the remaining pre-reaction and incubated at 30 °C for 15 minutes. Again, 12 μL of this reaction were removed and quenched. The remaining 88 μL mixture was then supplemented with additional 1 mM NADPH and adjusted to a final volume of 100 μL to prepare the AzeE pre-reaction. A portion of this mixture (12 μL) was removed and incubated with boiled AzeE as the negative control. AzeE (15 μM) was added to the remaining pre-reaction, the reaction adjusted to a final volume of 100 μL, and then incubated at 30 °C for 3 h. Twelve μL of the reaction were again quenched and kept aside. Another 12 μL of the reaction were removed and incubated with boiled AzeD as the negative control. AzeD (15 μM) was added to the remaining 76 μL of AzeE reaction in a final volume of 80 μL, and incubated at 30 °C for 3 h. A portion of this reaction (12 μL) was stopped using an equal volume of acetonitrile and stored for subsequent analysis. Finally, ferredoxin (10 μg), ferredoxin-$NADP^+$ reductase (0.1 U) and additional 1 mM NADPH were added into the AzeD reaction in a final volume of 100 μL, to prepare the AzeF pre-reaction. Twelve μL of this mixture were incubated with boiled AzeF as the negative control. In the meantime, AzeF (20 μM) was added into the remaining pre-reaction and the mixture incubated at 30 °C for 1 h, before being quenched using an equal volume of acetonitrile. All samples were stored at –20 °C for 30 min, and then centrifuged (20,000*g*, 10 min) to precipitate the proteins. The resulting supernatants were subjected to LC-HRMS analysis.

Attempt to identify the function of AzeG: AzeG (10 μM) was added to various stages of the reconstituted biosynthesis *in vitro* up to the AzeF reaction, and the products monitored using the same procedure as described above. However, no obvious changes in the product profile were detected.

With AZC as substrate: One-pot reactions followed the same procedure as that for Pro substrate until the AzeC step. In this case, AzeE (15 μM), AzeD (15 μM), AzeF (20 μM), ferredoxin (10 μg), ferredoxin-$NADP^+$ reductase (0.1 unit) and additional 1 mM NADPH were simultaneously added to the reaction mixture to a final volume of 100 μL, and incubated at 30 °C for 3 h. The results were compared to those from a negative control using boiled enzymes.

**Bioinformatic analysis of AzeD**

AzeD homologs were retrieved using BLAST against UniREF_90 database (E-value < 1e-10). Hmmsearch was then used with the PF00668.23 Condensation domain hmm profile as query to keep only homologs displaying a domain score > 50 and a length > 300. AzeB_$C_2$, AzeD and selected AzeD homologs were then aligned with the C domain sequence dataset from the study of

Wheadon and Townsend[9] using MAFFT v7.5 (with pairwise alignment option).[10] The resulting multiple alignment was then trimmed using Trimal v1.4.[11] SMS[12] and PhyML[13] were then used to infer the phylogenetic tree based on the best model according to the AIC criterion. Branch supports were calculated using transfer bootstrap method from 100 bootstrap replicates.[14] Genomic neighborhood of AzeD and its filtered homologs were obtained from EFI-GNT.[15]

**Production and purification of AzeD for X-ray crystallography**

Recombinant *E. coli* BL21(DE3) transformed with the expression plasmid was inoculated into 0.1 L of LB medium supplemented with kanamycin (50 µg/mL) and the cultures were incubated at 37 °C with agitation for 18 h. Several 2 L cultures of selective LB medium were seeded with 2 mL of a pre-culture and grown to an $OD_{600nm}$ of 0.6–0.7, and production of the recombinant proteins was induced by addition of 0.5 mM IPTG. Cultures were incubated at 20 °C with agitation for 24 h, and then the cells were harvested by centrifugation (4000 *g*, 4 °C, 30 min). Cell pellets were resuspended in 30 mL of lysis buffer (50 mM Tris-HCl, pH 7.8, 300 mM NaCl, 20 mM imidazole, 10% glycerol) and then the bacteria were lysed using four cycles of sonication on ice (15 kPsi, 4 °C, 2 min) (Sonifier S-250D, Branson Ultrasonics). Lysates were concentrated by centrifugation (48,000 g, 4 °C, 30 min) and the supernatant was filtered through a sterile membrane (0.22 µm) using a syringe (Merck).

Next, the supernatant was loaded onto a HisTrap affinity chromatography column ($Ni^{2+}$-sepharose, 10 mL) (GE Healthcare) equilibrated in buffer A (50 mM Tris-HCl pH 7.8, 300 mM NaCl, 20 mM imidazole, 10% glycerol) on an ÄKTA Avant system (Cytiva). The column was washed with 10 column volumes of lysis buffer at 5 mL/min, and then the protein eluted with buffer B containing 0.3 M imidazole at the same flow rate. Fractions containing the protein were concentrated using an Amicon Ultracel-10 concentrator (Merck). The sample was then injected through a loop (10 mL) onto a Superdex-75 size exclusion chromatography column (130 mL) (GE Healthcare) equilibrated in purification buffer (20 mM Tris-HCl, pH 7.8, 300 mM NaCl, 5% glycerol), using an ÄKTA Pure machine (Cytiva). The recombinant protein was eluted in the buffer at a flow rate of 1 mL/min, concentrated using an Amicon Ultracel-10 concentrator to 40 mg/mL, and then flash-frozen in liquid nitrogen and stored at −80 °C. The final sample concentration was measured using a NanoDrop 3000C spectrophotometer (Thermo Fisher Scientific), with extinction coefficients calculated using the ExPASy ProtParam tool.[16] Protein purity was assessed by 10% SDS-PAGE. Prior to crystallization trials, the monodispersity of AzeD dehydratase was verified by dynamic light scattering using a Zetasizer Nano-ZS (Malvern Panalytical).

**Crystallization of AzeD**

The crystallization conditions were screened using the sitting-drop vapor diffusion method in MRC plates (96 wells) (Grein Bio-One) with a Mosquito robot (SPT Labtech). Initial hits were obtained with the Index kit (Hampton Research), identifying the following condition: 100 mM bis-Tris methane, pH 6.5, 25% PEG-3350 (18 °C, 1–2 days of incubation). Crystals were manually reproduced using the hanging-drop vapor diffusion technique in Linbro® plates (24 wells) (Crystalgen), after mixing 1 µL of protein solution and 2 µL of crystallization buffer.

**X-ray diffraction, and construction/validation of the AzeD crystal structure**

Crystals were harvested with nylon cryoloops of suitable sizes (CrystalCap SPINE HT) (Hampton Research), then soaked in a cryoprotectant solution (mother liquor supplemented with 30% glycerol) for 10 s, and directly cryocooled in nitrogen gas flow at −180 °C. X-ray diffraction data were collected at the SOLEIL synchrotron (Saint-Aubin, France) on the PROXIMA-2A beamline. For this, crystals were successively mounted on a high-performance k-goniometer, then irradiated with a high-energy X-ray beam (12.7 keV (0.98 Å)) using the oscillation method over a standard angular range with an amplitude of 0.1°. Diffraction images were recorded using a single-photon counting EIGER X 9M detector (750 $image.s^{-1}$) (Dectris). The crystals belong to the P212121 space group (orthorhombic system) with the unit cell parameters a = 56.46 Å, b = 67.59 Å, c = 116.74 Å, a = β = γ = 90°. Datasets were indexed and integrated using the XDS software,[17] and scaled and reduced to the asymmetric unit using the aimless program (CCP4).[18,19] Their quality was assessed based on relevant parameters such as maximum resolution, signal-to-noise ratio, completeness, multiplicity and correlation coefficient CC1/2 (**Table S5**).

Initial phases were determined by molecular replacement using the MOLREP program[20] based on a structural model generated using the AlphaFold2 software.[21] The structure was constructed and refined by iterative cycles of manual reconstruction in the electron density maps using the WinCoot program,[22] followed by restrained refinement at 1.55 Å using the REFMAC5 program[23] (**Table S5**). The final model geometry was validated using the MolProbity program[24] and the wwPDB OneDep server.[25] The structure of the AzeD dehydratase has been deposited in the PDB database under the accession code 8S6L. Binary complex structures with the substrate were subsequently modeled by molecular docking using the CB-Dock2 program.[26] The relevant **Figures 5** and **S27-S28** were prepared using the PyMOL program.[27]

**Small-angle X-ray scattering (SAXS) analysis of AzeD**

Size-exclusion chromatography-small angle X-ray scattering (SEC-SAXS) data were acquired on the SWING beamline at the SOLEIL synchrotron (Saint Aubin, France). Frame recording was carried out using an EIGER X 4M detector at an energy of 12 keV. The sample/detector distance was set to 2000 mm, giving rise to scattering vectors $q$ ranging from 0.0005–0.55 $Å^{-1}$. The protein samples were injected using an automatic online sample changer[28] into a pre-equilibrated size-exclusion chromatography column (Bio-SEC 150, Agilent) coupled to an HPLC, at a temperature of 15 °C. After equilibrating the column in the protein buffer (20 mM Tris-HCl, pH 7.8, 300 mM NaCl, 5% glycerol), 50 µL of protein sample at 40 mg/mL was injected. Data corresponding to buffer background and the eluted protein were recorded over 600 successive frames. The protein concentration downstream of the elution column was monitored *in situ* via the absorbance at 280 nm using a spectrophotometer.

The in-house application FOXTROT[29] was used to perform initial data treatment (reduction to absolute units, frame averaging, and solvent subtraction). FOXTROT was then employed to determine the $R_g$ (radius of gyration), and the $I(0)$ value of each acquisition frame. We observed an essentially constant $R_g$ for the majority of the concentrations present in the gel filtration peaks, showing that the measurements were concentration-independent. The frames exhibiting a constant $R_g$ as a function of $I(0)$ were then averaged. The distance distribution function $P(r)$ and the maximum particle diameter $D_{max}$ were also calculated using GNOM.[30] The molecular weight of AzeD was determined from the acquired SAXS data using PRIMUS.[31] The SAXS data are presented in **Table S6**.

**Computational docking of** 5 **into the AzeD active site**

Binary complex structures with the substrate **5** were modeled by molecular docking using the online program CB-Dock2.[32] The average energy value associated with **5** binding at the active site was $-8.5 \pm 0.1$ $kcal \cdot mol^{-1}$ (5 calculations).

**Circular dichroism measurements**

Circular dichroism spectra were recorded with 5 μM each of the wild-type AzeD and mutants in 50 mM Tris-HCl (pH 8.0) on a Jasco J-810 spectropolarimeter. Mean residue ellipticity was obtained using the formular: Θ/(10*L*N*c) (Θ, observed ellipticity; L, light path (cm); N, number of peptide bonds; c, protein concentration (M)). The graphs were generated using the LabPlot tool.

**Table S1**. Strains and plasmids used in this study.

| | Characteristics/Function | Source |
|---|---|---|
| Strains | | |
| *P. aeruginosa* AB16.2 | Colistin-resistant mutant of PAO1 strain; azetidomonamide production | [3] |
| *P. aeruginosa* AB16.2ΔazeA | AB16.2 with in-frame deletion of 189 bp in PA3326; azetidomonamide production | [2] |
| *P. aeruginosa* AB16.2ΔazeA_ΔazeD | AB16.2Δ*azeA* with in-frame deletion of 621 bp in PA3329 | This study |
| *P. aeruginosa* AB16.2ΔazeA_ ΔazeE | AB16.2Δ*azeA* with in-frame deletion of 366 bp in PA3330 | This study |
| *P. aeruginosa* AB16.2ΔazeA_ ΔazeF | AB16.2Δ*azeA* with in-frame deletion of 444 bp in PA3331 | This study |
| *P. aeruginosa* AB16.2ΔazeA_ ΔazeG | AB16.2Δ*azeA* with in-frame deletion of 225 bp in PA3332 | This study |
| *E. coli* DH5α | $F^-$ φ80*lac*ZΔM15 Δ(*lac*ZYA-*arg*F)U169 *rec*A1 *end*A1 *hsd*R17($r_K^-$, $m_K^+$) *pho*A *sup*E44 $\lambda^-$*thi*-1 *gyr*A96 *rel*A1; plasmid propagation | Thermo Fischer Scientific ™ |
| CC118λ*pir* | Δ(*ara-leu*) *ara*D Δ*lac*X74 *gal*E *gal*K *pho*A20 *thi*-1 *rps*E *rpo*B *arg*E(Am) *rec*Al lysogenized with λ*pir* phage | [33] |
| HB101 | *sup*E44 *hsd*S20($r_B^-$, $m_B^-$) *rec*A13 *ara*-14 *pro*A2 *lac*Y1 *gal*K2 *rps*L20 *xyl*-5 *mtl*-1 *leu*B6 *thi*-1 | [34] |
| *E. coli* BL21(DE3) | F– *ompT gal dcm lon hsdSB*(*rB–mB–*) λ(DE3 [*lacI lacUV5-T7p07 ind1 sam7 nin5*]) [*malB*+]K-12(λS); protein expression | Novagen |
| *E. coli* BAP1 | BL21(DE3) ΔprpRBCD::T7prom－sfp,T7prom－prpE； protein expression | [7] |
| **Plasmids for protein expression** | | |
| pET28a(+) | Protein expression vector | Novagen |
| pET29b(+) | Protein expression vector | Novagen |
| pBG106 | Protein expression vector | Center for Structural Biology, Vanderbilt University |

| | | |
|---|---|---|
| pET28-azeB | *azeB* inserted between *Nde*I/*Hind*III in pET28a(+); expression of N-$His_6$-AzeB | This study |
| pET28-azeB_A1L | Expression of N-$His_6$-tagged, truncated AzeB (AA 1–1046) spanning the C1-A1-PCP1 domains | [3] |
| pET28-azeB_A1LC2 | Expression of N-$His_6$-tagged, truncated AzeB (AA 1–1479) spanning the C1-A1-PCP1-C2 domains | This study |
| pET28-azeB_A2LTE | Expression of N-$His_6$-tagged, truncated AzeB (AA 1024–2353) spanning the C2-A2-PCP2-TE domains | This study |
| pBG106-azeB_C2A2 | Expression of N-$His_{10}$-tagged, truncated AzeB (AA 1033–1981) spanning the C2-A2 domains | This study |
| pET28-azeC | *azeC* inserted between *Nde*I/*Hind*III in pET28a(+); expression of N-$His_6$-AzeC | This study |
| pET28-azeD | *azeD* inserted between *Nde*I/*Hind*III in pET28a(+); expression of N-$His_6$-AzeD | This study |
| pET28-azeD-st | *azeD* inserted between *Bam*HI/*Hind*III in pET28a(+); expression of N-$His_6$-AzeD for crystallization | This study |
| pET28-azeD-H130A | Expression of N-$His_6$-AzeD single mutant H130A | This study |
| pET28-azeD-T290A | Expression of N-$His_6$-AzeD single mutant T290A | This study |
| pET28-azeD-C355A | Expression of N-$His_6$-AzeD single mutant C355A | This study |
| pET28-azeD-S357A | Expression of N-$His_6$-AzeD single mutant S357A | This study |
| pET29-azeE | *azeE* lacking a stop codon inserted between *Nde*I/*Xho*I in pET29b(+); expression of C-$His_6$-AzeE | This study |
| pET28-azeF | *azeF* inserted between *Nde*I/*Hind*III in pET28a(+); expression of N-$His_6$-AzeF | This study |
| **Plasmids for gene inactivation in *P. aeruginosa*** | | |
| pRK2013 | Helper plasmid for mobilization of non-self-transmissible plasmids; ColE1 $Tra^+$ $Mob^+$ $Kan^r$ | [6] |
| pKNG101 | Marker exchange suicide vector in *P. aeruginosa*; *sac*BR *mob*RK2 *ori*R6K $Str^r$ | [5] |
| pKNGΔ3326 | *BamH*I/*Apa*I 1.134-kb fragment composed of sequences flanking the 5′ and 3′ ends of *azeA*, cloned in pKNG101 | This study |
| pKNGΔ3329 | *BamH*I/*Apa*I 1.063-kb fragment composed of sequences flanking the 5′ and 3′ ends of *azeD*, cloned in pKNG101 | This study |
| pKNGΔ3330 | *BamH*I/*Apa*I 1.072-kb fragment composed of sequences flanking the 5′ and 3′ ends of *azeE*, cloned in pKNG101 | This study |
| pKNGΔ3331 | *BamH*I/*Apa*I 1.006-kb fragment composed of sequences flanking the 5′ and 3′ ends of *azeF*, cloned in pKNG101 | This study |
| pKNGΔ3332 | *BamH*I/*Apa*I 1.067-kb fragment composed of sequences flanking the 5′ and 3′ ends of *azeG*, cloned in pKNG101 | This study |

$Tc^r$, tetracycline resistance; $Str^r$, streptomycin resistance; $Kan^r$, kanamycin resistance; $Zeo^r$, zeocin resistance.

**Table S2.** Primers used in this study.

| **Primer name** | **Sequence (5′→3′)[a]** |
|---|---|
| **For construction of protein expression plasmids** | |
| PA004_3327F | GATATGCATATGGTTCGTTTCGCTCGCTTG |
| PA005_3327R | GATATGAAGCTTTCATGGGCGGGTCCGTTCG |
| XH008_A1LC2R | ATACTAAAGCTTTCAGCTGACGACGTCGACCAG |
| PA013_A2LTEF | GATATGCATATGCTCGCGGAGCGAAGCTACC |
| pBG106-C2_A2_Fw | TTTCCGGATCCATGGAACTGGTTTCCGAAGTCGACAAGC |
| pBG106-C2_A2_Rv | TTTCCAAGCTTTTACTCGATGTTCGTCCCGTGCTCCAG |
| SN30_AzeC _F | GGAATTCCATATGAGCAGACATCCCCTGAA |
| SN30_AzeC _R | CCCAAGCTTTCATTCGACTCCTGGGAAG |
| SN55_AzeD_F | GGAATTCCATATGGCCGAGATACGACGC |
| SN56_AzeD_R | CCCAAGCTTTCATTGCTCCTCGAGGGC |
| AzeD-F_crystallo | TTTCCGGATCCATGGCCGAGATACGACGC |
| AzeD-R_crystallo | TTTCCAAGCTTTTATTGCTCCTCGAGGGCGGATC |
| SN57_AzeE_F | GGAATTCCATATGTTATTCACCAGCAAACC |
| SN63_AzeE_R | CCGCTCGAGGGCACGTTGCTCCTCCTT |
| SN61_AzeG_F | GGAATTCCATATGAATGCCAAGGAAATTC |
| SN62_AzeG_R | CCCAAGCTTTCAGGCTCCCTGGACGAT |
| SN59_AzeF_F | GGAATTCCATATGCCTGATCGCAAACTGA |
| SN60_AzeF_R | CCCAAGCTTTCATTGGGCGATCCTCTCG |
| SN89_AzeD_H130A_F | GGTACCGGCCATCATCGCCGACGGCACTAC |
| SN90_AzeD_H130A_R | ATGATGGCCGGTACCACGACCAGCAGGTCG |
| SN95_AzeD_T290A_F | CGTCGCCGCGGTACCGGTGGTGCTGGACAT |
| SN96_AzeD_T290A_R | GGTACCGCGGCGACGTAGGTGCCTACTTCG |
| SN97_AzeD_C355A_F | CAACCTGGCCTCCTCCAACATCGGTCGCTA |
| SN98_AzeD_C355A_R | GAGGAGGCCAGGTTGATCGGCCCTTCGGCT |
| SN99_AzeD_S357A_F | GTGCTCCGCCAACATCGGTCGCTATCCGTT |
| SN100_AzeD_S357A_R | ATGTTGGCGGAGCACAGGTTGATCGGCCCT |
| **For construction of *P. aeruginosa* mutants** | |
| PCRi PA3326 A1 | CGCCGGTCTTTTCGAAGTTG |
| PCRi PA3326 A2 | GCCGTACCGGGTATTGGGCAGCGAATA |
| PCRi PA3326 A3 | ATACCCGGTACGGCCTGGTCAACAAGA |
| PCRi PA3326 A4 | CGCCCGTTACTTCAATCCCT |
| PCRi PA3328 A1 | GAACCCTATGTCGAGACGGTC |
| PCRi PA3328 A2 | ACGTTGCCCAGTTCGATGCGGCAGTC |
| PCRi PA3328 A3 | GAACTGGGCAACGTCTATTGGTGGGGC |
| PCRi PA3328 A4 | TTCGACTCCTGGGAAGGTCA |
| PCRi PA3329 A1 | AACGAACTGGCACTGACCTT |
| PCRi PA3329 A2 | AGGTTGAATTGTTCGGCGAGGGTCAG |

| | |
|---|---|
| PCRi PA3329 A3 | GAACAATTCAACCTGTGCTCCTCCAAC |
| PCRi PA3329 A4 | CCAATGGATTCCTCGTCGGT |
| PCRi PA3330 A1 | CGCCATCAACTCCAGCCAT |
| PCRi PA3330 A2 | GGTATCGCAGATCCCGGCGTTGTTCA |
| PCRi PA3330 A3 | GGATCTGCGATACCTACCTGCGCTTCC |
| PCRi PA3330 A4 | TCGAGGTTGAGCATGTGGTG |
| PCRi PA3331 A1 | ACCAAGCCCGCCGATGTC |
| PCRi PA3331 A2 | CGAAGCCCATGTGGTGGCTGAGTCCC |
| PCRi PA3331 A3 | CCACATGGGCTTCGAGACCACCATGAA |
| PCRi PA3331 A4 | GGAGGAAGGTGATCGGCAG |
| PCRi PA3332 A1 | GCGAATACATCCTGGTCTCCA |
| PCRi PA3332 A2 | AGAGCAGGGCGTAGGGGAATTCGAGTAC |
| PCRi PA3332 A3 | CTACGCCCTGCTCTACCGCGACTTCTG |
| PCRi PA3332 A4 | CTGAAAGTGTCCACCCCGAC |

[a]: Restriction sites are underlined.

**Table S3**. Summary of azetidomonamides and pathway intermediates detected in extracts *in vivo* or in enzyme assays *in vitro*.

| Structure | Retention time (min) | Presence *in vivo* | Produced *in vitro* |
|---|---|---|---|
| 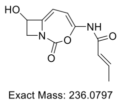 **1** | 5.4 | ✓ | ✓ |
| 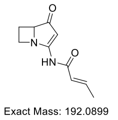 **2** | 4.9 | ✓ | ✓ |
| 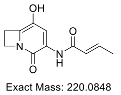 **3** | 5.0 | ✓ | ✓ |
| 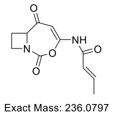 **4** | 5.9 | ✓ | ✓ |
| 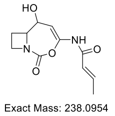 **5** | n.a. | n.d. | n.d. |
| 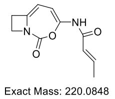 **6** | 6.4 | ✓ | ✓ |
| 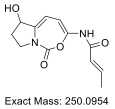 **1′** | 5.6 | ✓ | ✓ |
| 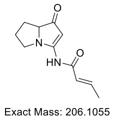 **2′** | 5.3 | ✓ | ✓ |
| 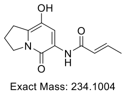 **3′** | 5.5 | ✓ | ✓ |

| | | | |
|---|---|---|---|
| 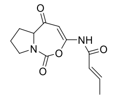 Exact Mass: 250.0954 **4′** | 6.4 | ✓ | ✓ |
| 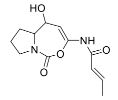 Exact Mass: 252.1110 **5′** | 4.0 | n.d | ✓ |
| 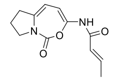 Exact Mass: 234.1004 **6′** | 7.1 | ✓ | ✓ |

n.a. not available; n.d. not detected

**Table S4.** Quantitative comparison of *aze* pathway products and intermediates detected in *P. aeruginosa* mutants. Quantification was based on the calculated areas of the corresponding peaks in the extracted ion chromatograms. For each compound, the most intense peak was taken as 100%.

| Compound | Parental strain | Δ*azeD* | Δ*azeE* | Δ*azeF* | Δ*azeG* |
|---|---|---|---|---|---|
| **1** | 100% | n.d. | n.d. | n.d. | n.d. |
| **2** | 92% | 69% | 100% | 28% | 24% |
| **3** | 100% | n.d. | n.d. | n.d. | n.d. |
| **4** | n.d. | 55% | 100% | 21% | n.d. |
| **6** | 3% | n.d. | n.d. | 100% | n.d. |
| **1′** | 100% | n.d. | n.d. | n.d. | n.d. |
| **2′** | 100% | 9.5% | 19% | 9.2% | 6.0% |
| **3′** | 100% | 4.0% | 3.3% | 2.8% | n.d. |
| **4′** | 100% | n.d. | n.d. | n.d. | n.d. |
| **6′** | 9.4% | n.d. | n.d. | 100% | n.d. |

n.d. not detected

**Table S5.** Recording, processing and refinement of the crystallographic data acquired on the AzeD dehydratase.

| **Recording and processing data** | **AzeD** |
|---|---|
| Space group | $P2_12_12_1$ |
| Cell dimensions | |
| a, b, c [Å] | 56.46, 67.59, 116.68 |
| α, β, γ [°] | 90.00, 90.00, 90.00 |
| Wavelength [Å] | 0.980112 |
| Resolution [Å] | 44.20–1.55 (1.59–1.55) |
| *R*merge [%] | 6.4 (107.9) |
| *R*pim [%] | 1.9 (30.3) |
| $I / \sigma I$ | 20.0 (1.9) |
| Number of total observations | 857500 |
| Number of unique reflections | 65206 |
| Completeness [%] | 99.9 (98.7) |
| Multiplicity | 13.1 (13.6) |
| Wilson *B*-factors [$Å^2$] | 31.535 |
| Correlation coefficient $CC_{1/2}$ | 99.9 (76.9) |
| **Refinement data** | |
| Resolution [Å] | 44.20–1.55 |
| Number of reflections | 61945 |
| *R*, *R*free | 0.1457, 0.1945 |
| Number of atoms | 3763 |
| Protein | 3411 |
| Other ligands/ions | 12 |
| Water | 340 |
| Average *B*-factors, all atoms [$Å^2$] | 24.346 |
| R.m.s deviations | |
| Bond lengths [Å] | 0.0080 |
| Bond angles [°] | 1.4670 |
| **PDB ID** | **8S6L** |

**Table S6.** Recording and processing of the SAXS data acquired on the AzeD dehydratase.

| **Recording and processing data** | **AzeD** |
|---|---|
| Guinier curve analysis | |
| $I(0)$ [$cm^{-1}$] | 0.110 ± $8.0.10^{-5}$ |
| $R_g$ [Å] | 24.13 ± 0.03 |
| $q_{min}$ [$Å^{-1}$] | 0.034 |
| $q.R_g$ max | 1.08 |
| Correlation coefficient $R^2$ | 0.97 |
| $P(r)$ function analysis | |
| $q$ range [$Å^{-1}$] | 0.01− 0.32 |
| $I(0)$ [$cm^{-1}$] | 0.110 |
| $R_g$ \|Å] | 24.52 |
| $D_{max}$ [Å] | 91.63 |
| Correlation coefficient $R^2$ | 0.83 |
| Molecular weight analysis [kDa] | |
| Porod invariant ($Q_p$) | 43.71 |
| MoW | 49.80 |
| Correlation volume ($V$c) | 44.10 |
| Bayesian inference | 44.73 |
| Absolute scale | n.d. |

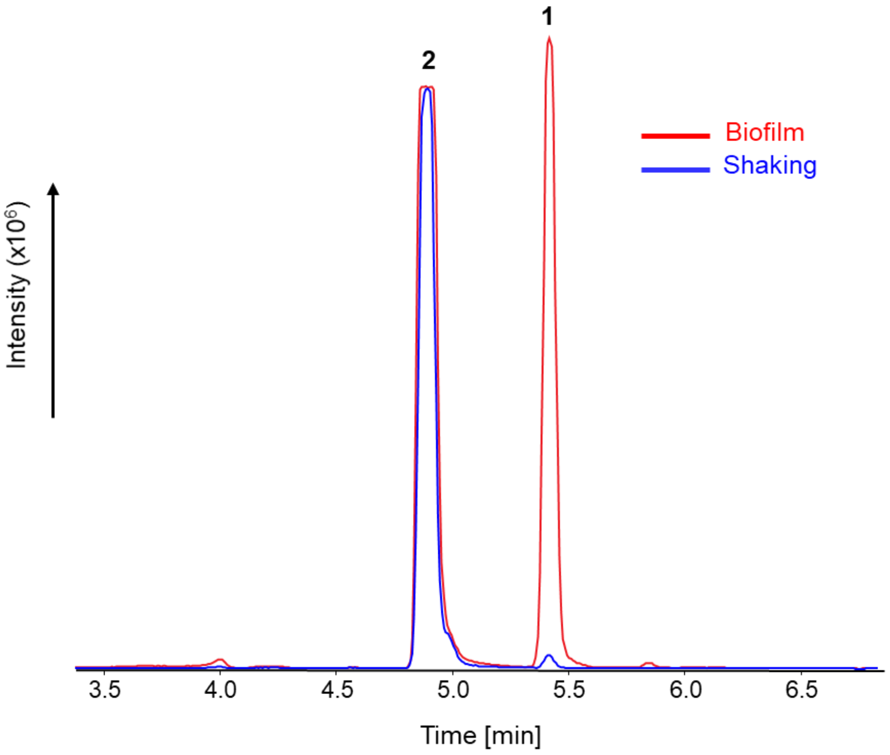


**Figure S1.** Comparison of production levels of azetidomonamides in static (biofilm) and shaking conditions by LC-HRMS. Combined EICs of $[M+H]^+$ 237.0875 and 193.0977 corresponding to azetidomonamide A **1** and B **2**, respectively, are shown. Ten mL of *P. aeruginosa* AB16.2 strain were cultured in 2×M9 medium without shaking (red trace) and with shaking (180 rpm; blue trace) in a 100 mL flask at 30 °C for 48 h. The biofilm condition dramatically increased the production of azetidomonamide A **1**.

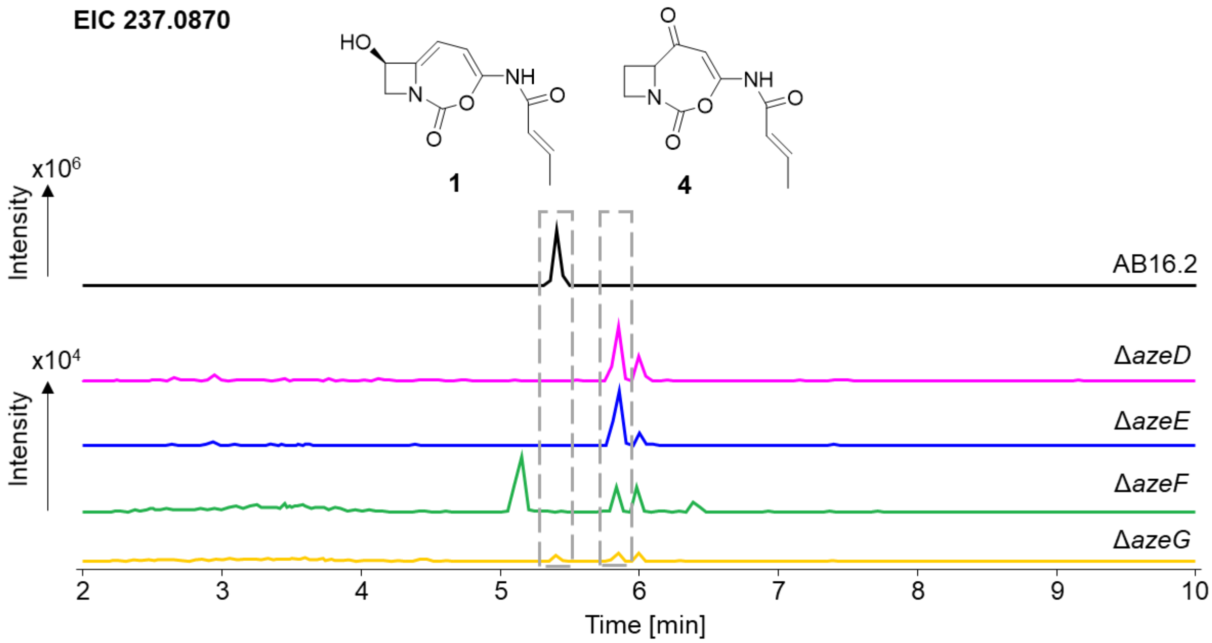


**Figure S2.** Accumulation of low amounts of intermediate **4** in the mutant strains. EICs of *m/z* 237.0870 corresponding to the $[M+H]^+$ ion of **1** and **4** are shown. For comparison, no **4** could be detected in the parental AB16.2 strain. A quantitative comparison of the yields is provided in **Table S4**.

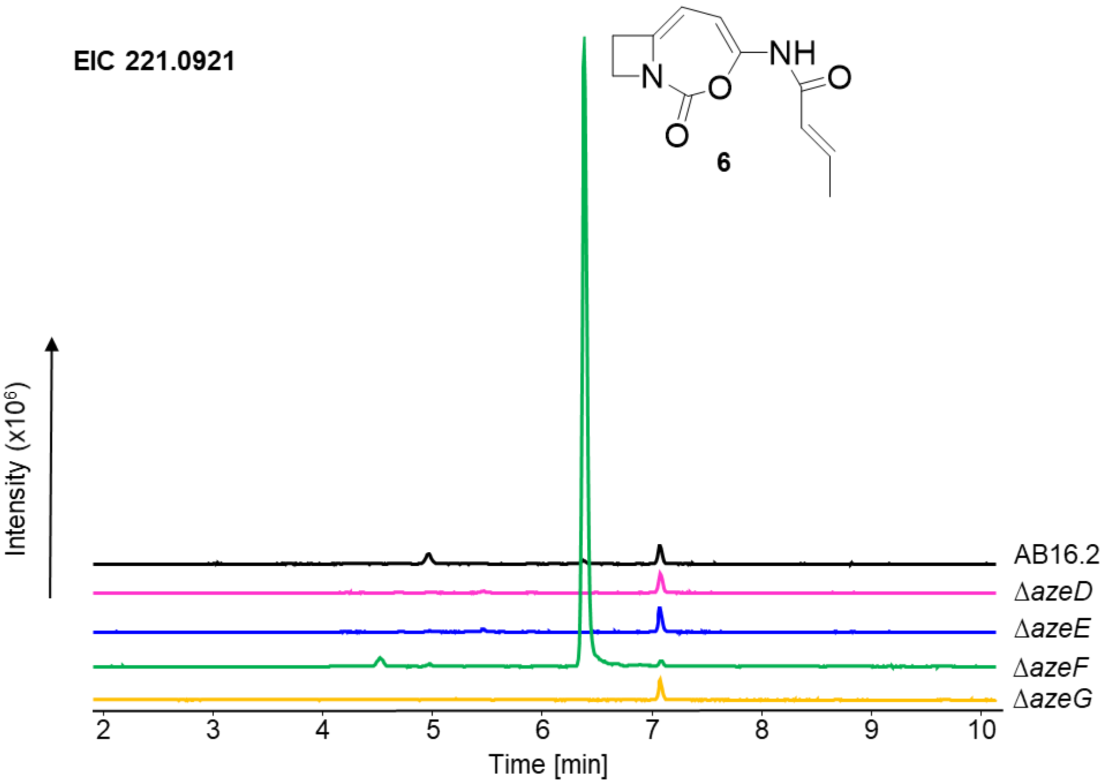


**Figure S3.** Accumulation of azetidomonamide C **6** in the Δ*azeF* mutant strain. EICs of *m/z* 221.0921 corresponding to the $[M+H]^+$ ion of **6** are shown. A quantitative comparison of the yields is provided in **Table S4**.

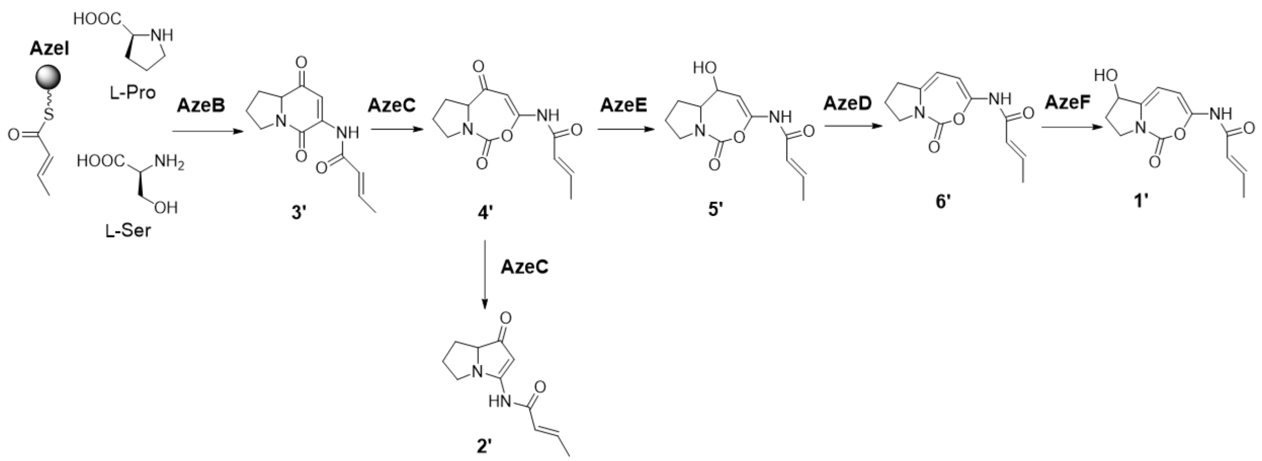


**Figure S4.** Proposed biosynthesis of Pro-derived compounds by the *aze* pathway.

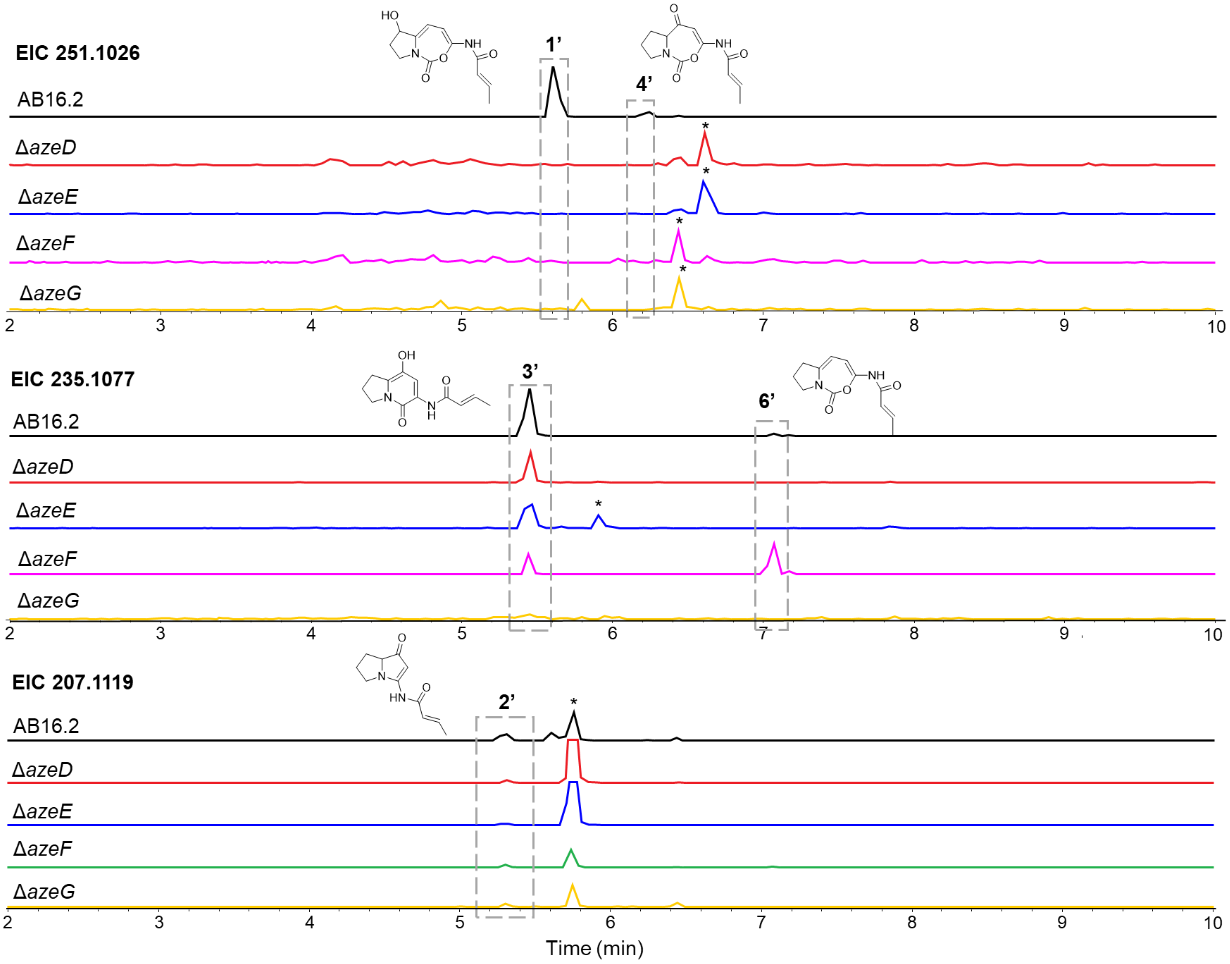


**Figure S5.** Detection of Pro-derived intermediates in AB16.2 and its single deletion mutant strains by LC-HRMS. EICs of the target compounds are shown. * denotes unrelated peaks with the same mass. A quantitative comparison of the yields is provided in **Table S4**.

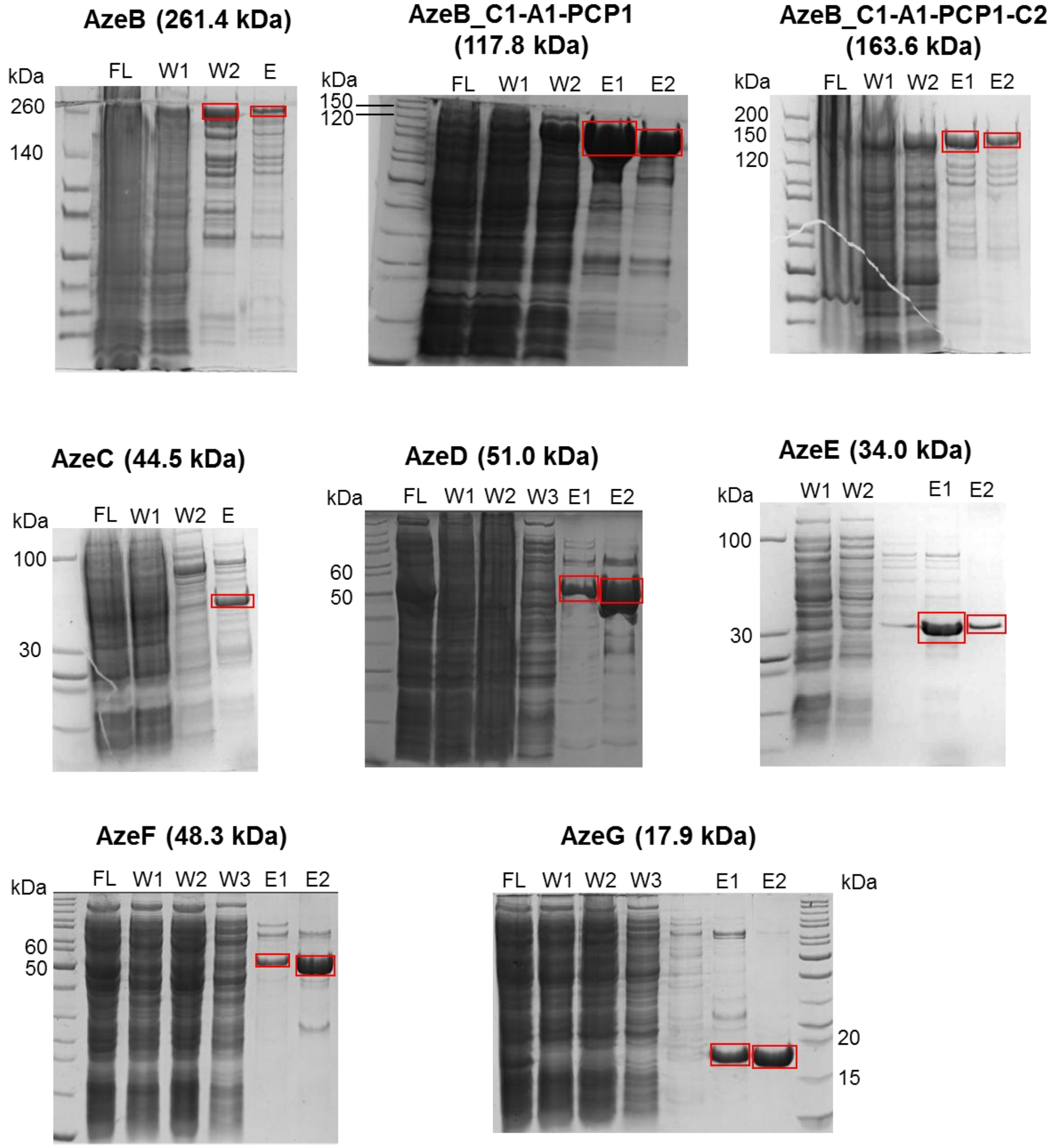


**Figure S6:** SDS-PAGE analysis of recombinant proteins purified by nickel affinity chromatography, and which were employed in the *in vitro* assays. FL, flowthrough; W, wash; E, elution. Targeted proteins in the elution fractions are boxed.

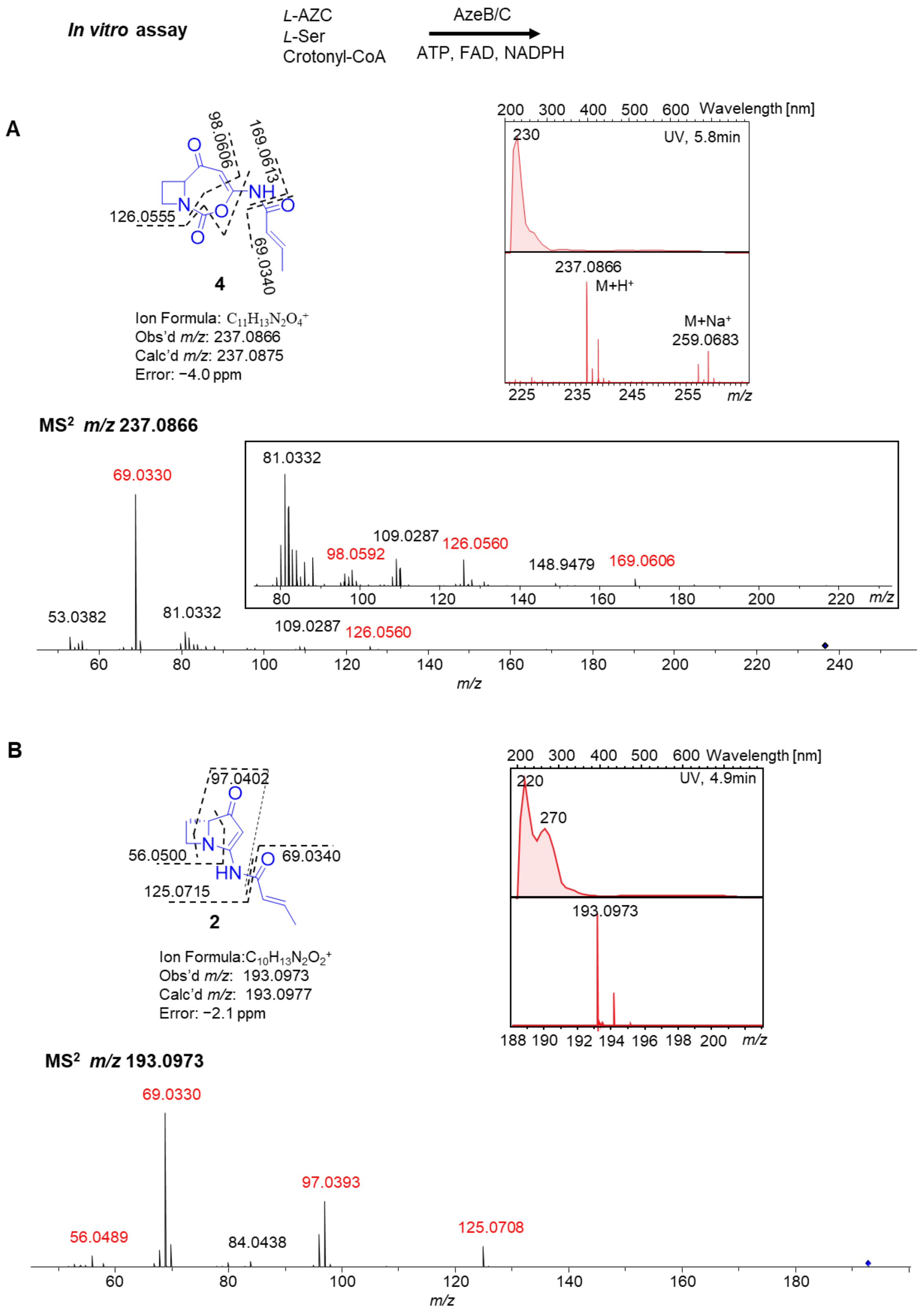


**Figure S7.** Characterization of products in AzeB/AzeC assays *in vitro* using *L*-AZC as substrate. The structures, UV, MS, $MS^2$ spectra and fragment annotation of **4** (panel **A**) and **2** (panel **B**) are shown.

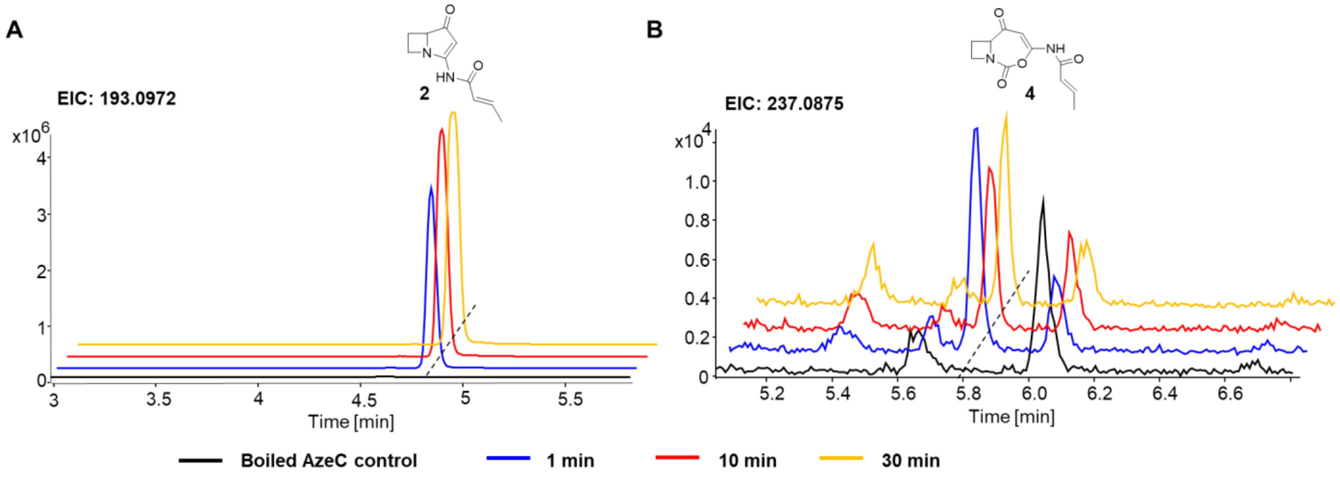


**Figure S8.** Time-course of AzeB/AzeC reaction with *L*-AZC as substrate *in vitro*. EICs of the corresponding $[M+H]^+$ ions are shown.

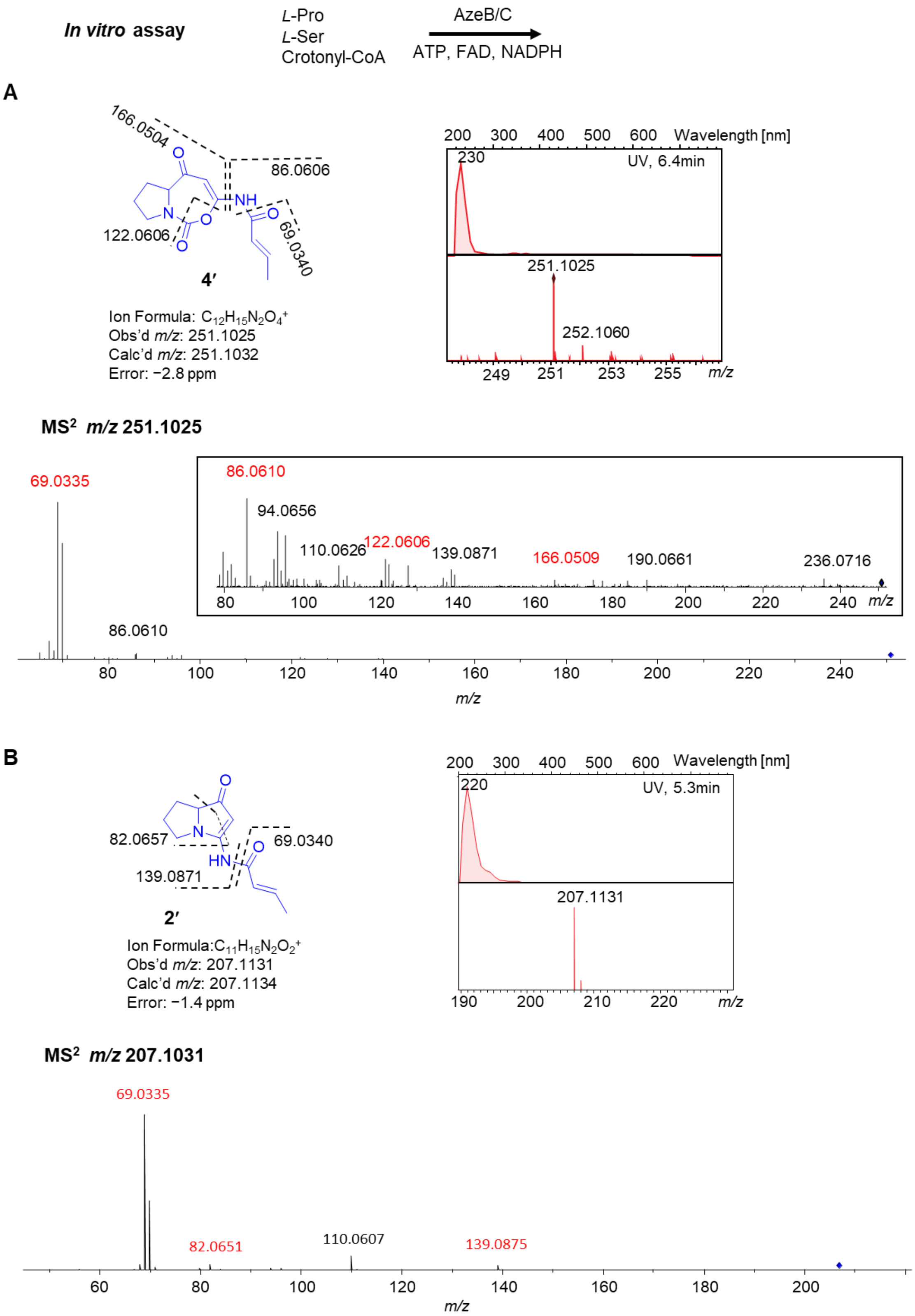


**Figure S9.** Characterization of products in AzeB/AzeC assays *in vitro* using Pro as substrate. The structures, UV, MS, MS$^2$ spectra and fragment annotation of **4′** (panel **A**) and **2′** (panel **B**) are shown.

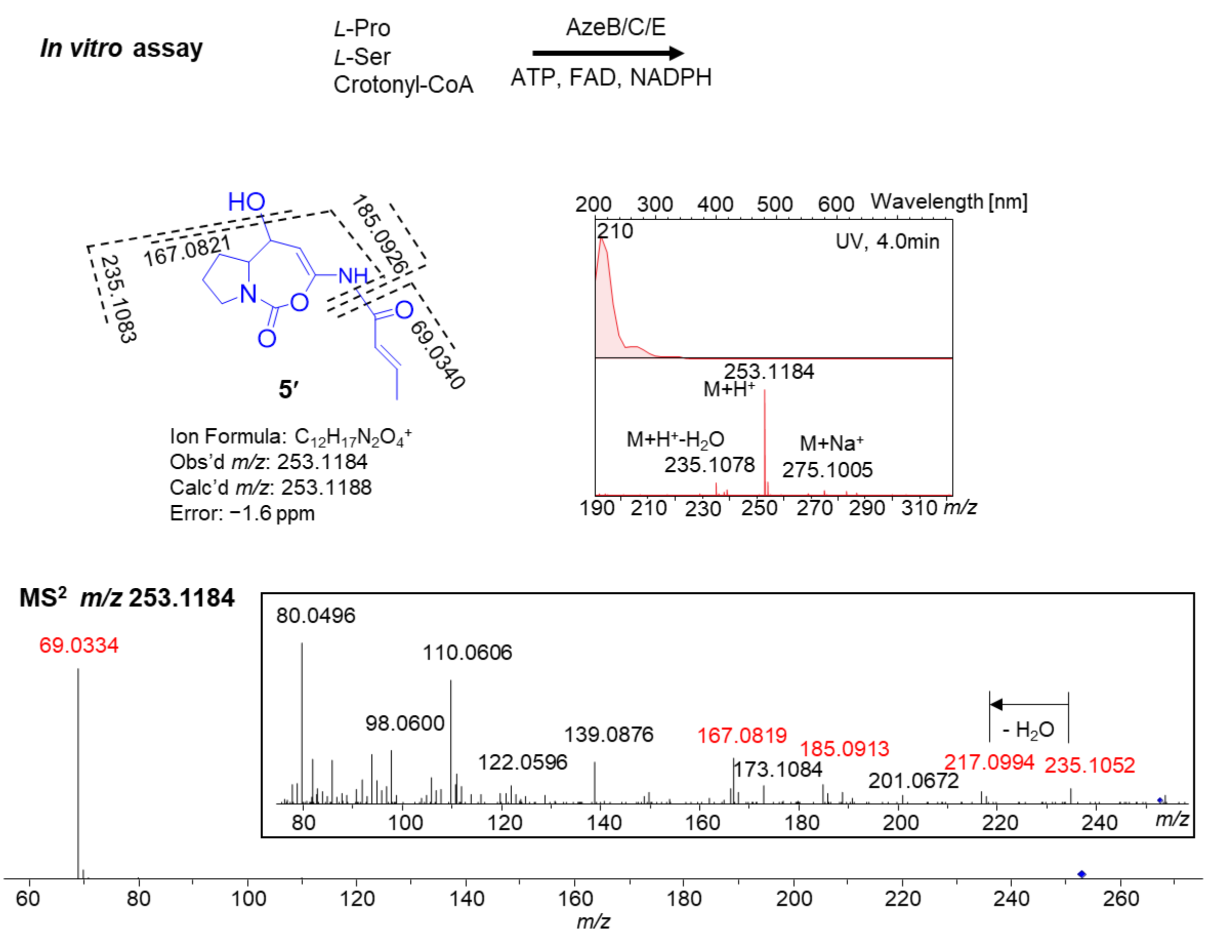


**Figure S10.** Characterization of AzeE *in vitro* using Pro as substrate. The structure, UV, MS, MS$^2$ spectra and fragment annotation of the reaction product **5′** are shown.

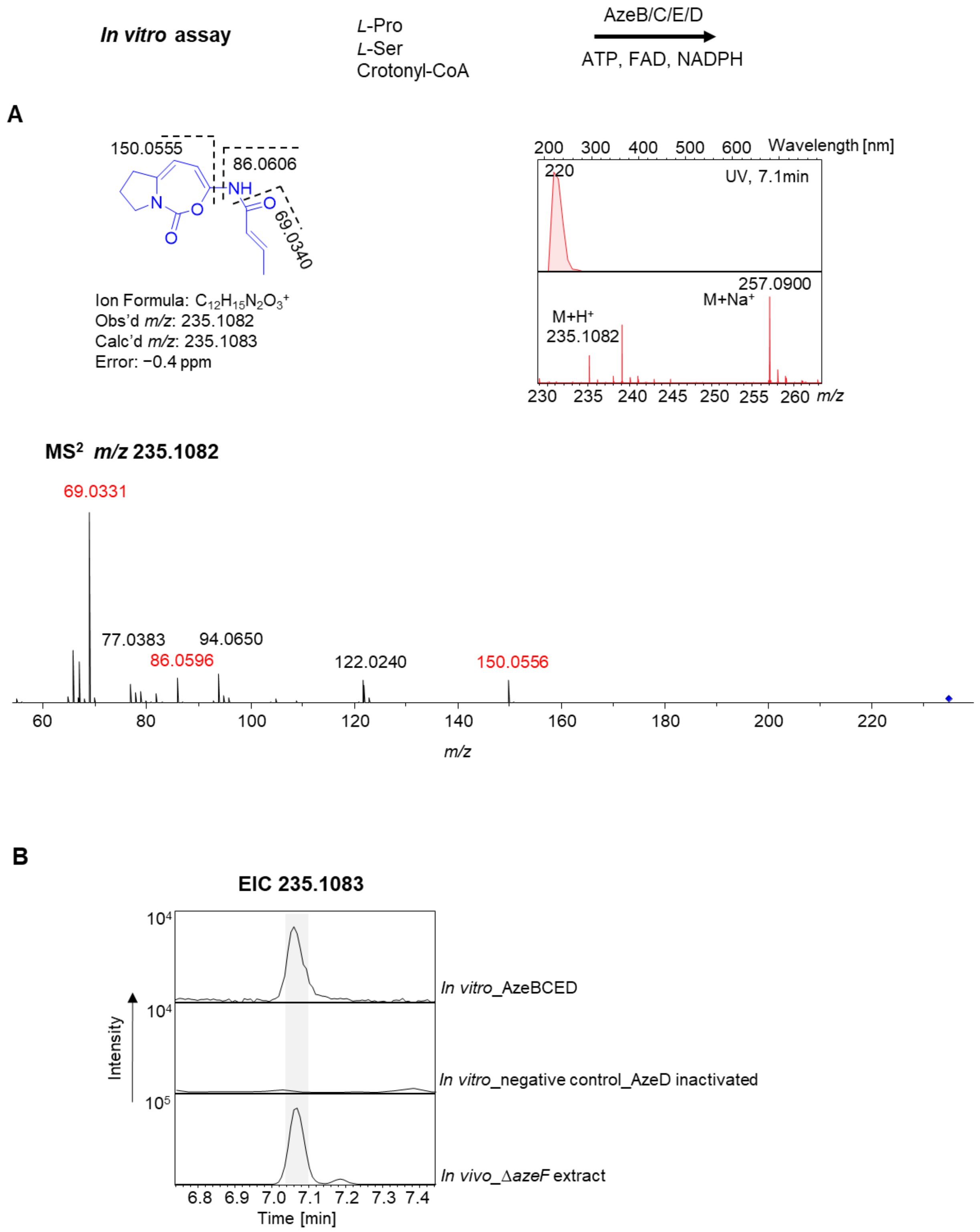


**Figure S11.** Characterization of AzeD *in vitro* using Pro as substrate. **A**) The structure, UV, MS, $MS^2$ spectra and fragment annotation of the reaction product **6′**. **B**) Comparison of EICs corresponding to $[M+H]^+$ ion of **6′** produced *in vivo* and *in vitro*.

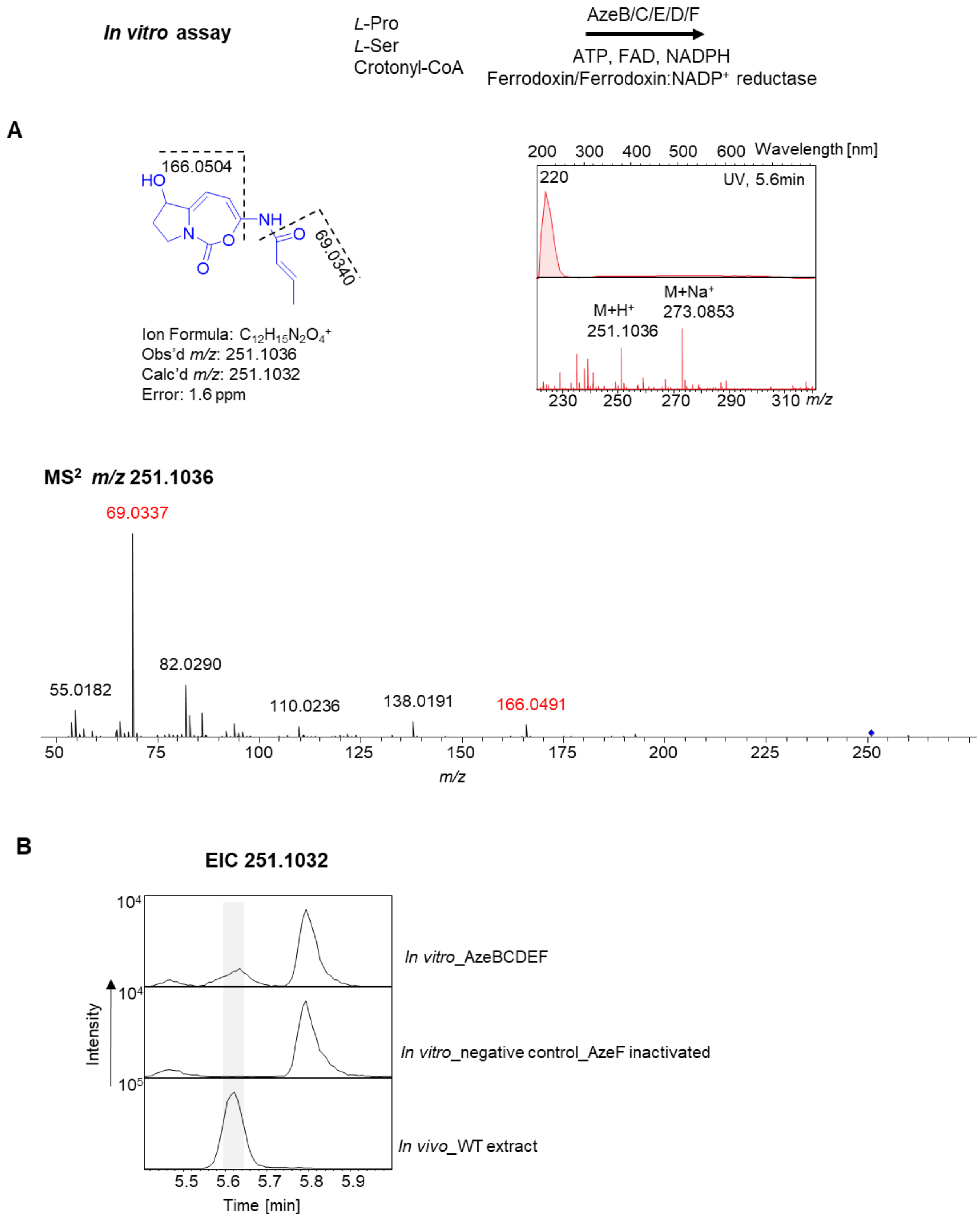


**Figure S12. Total biosynthesis *in vitro* of Pro-incorporated 1′.** **A**) The structure, UV, MS, $MS^2$ spectra and fragment annotation of **1′**. **B**) Comparison of EICs corresponding to $[M+H]^+$ ion of **1′** produced *in vivo* and *in vitro*.

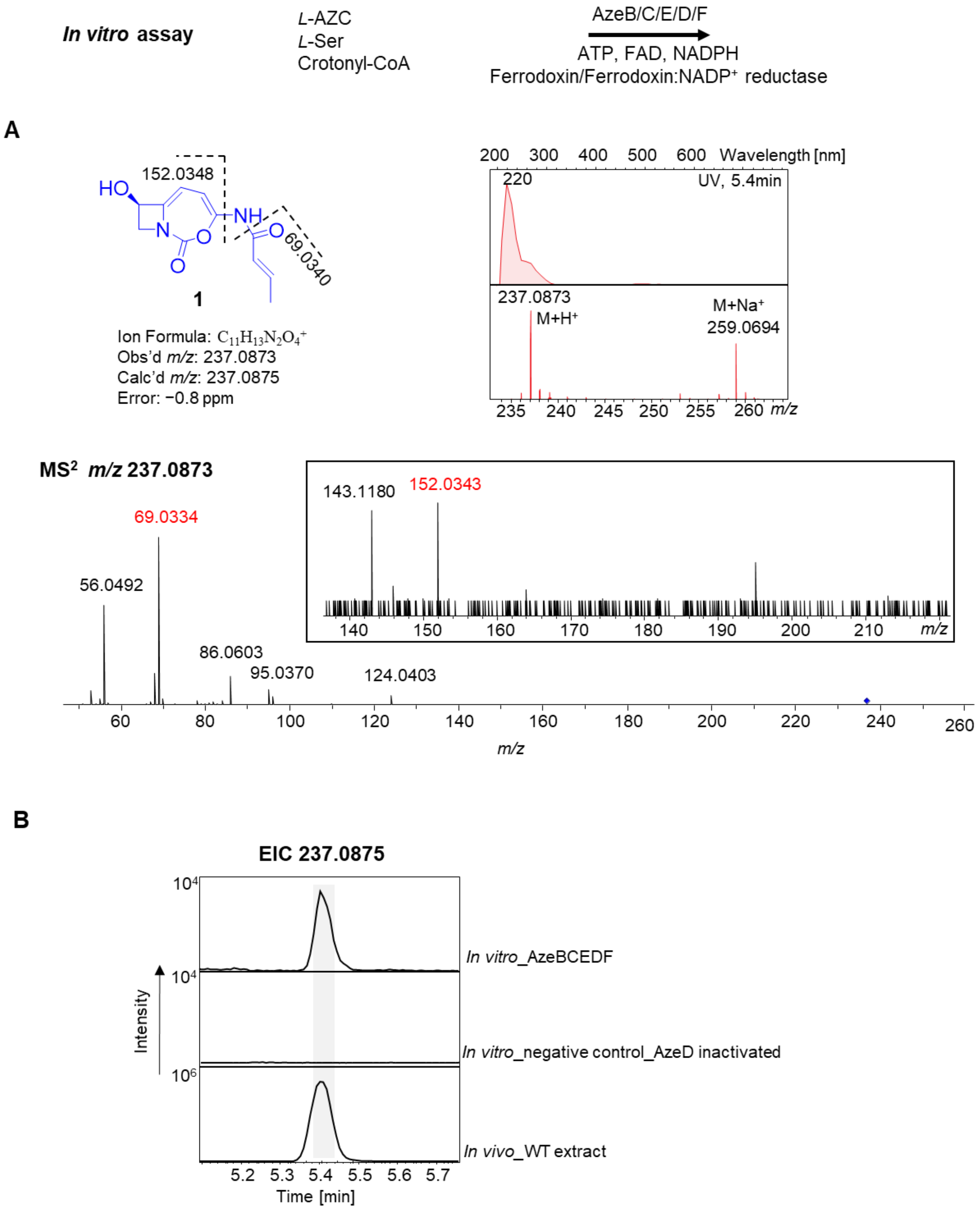


**Figure S13.** Total biosynthesis *in vitro* of azetidomonamide A **1**. **A**) The structure, UV, MS, MS$^2$ spectra and fragment annotation of **1**. **B**) Comparison of EICs corresponding to $[M+H]^+$ ion of **1** produced *in vivo* and *in vitro*.

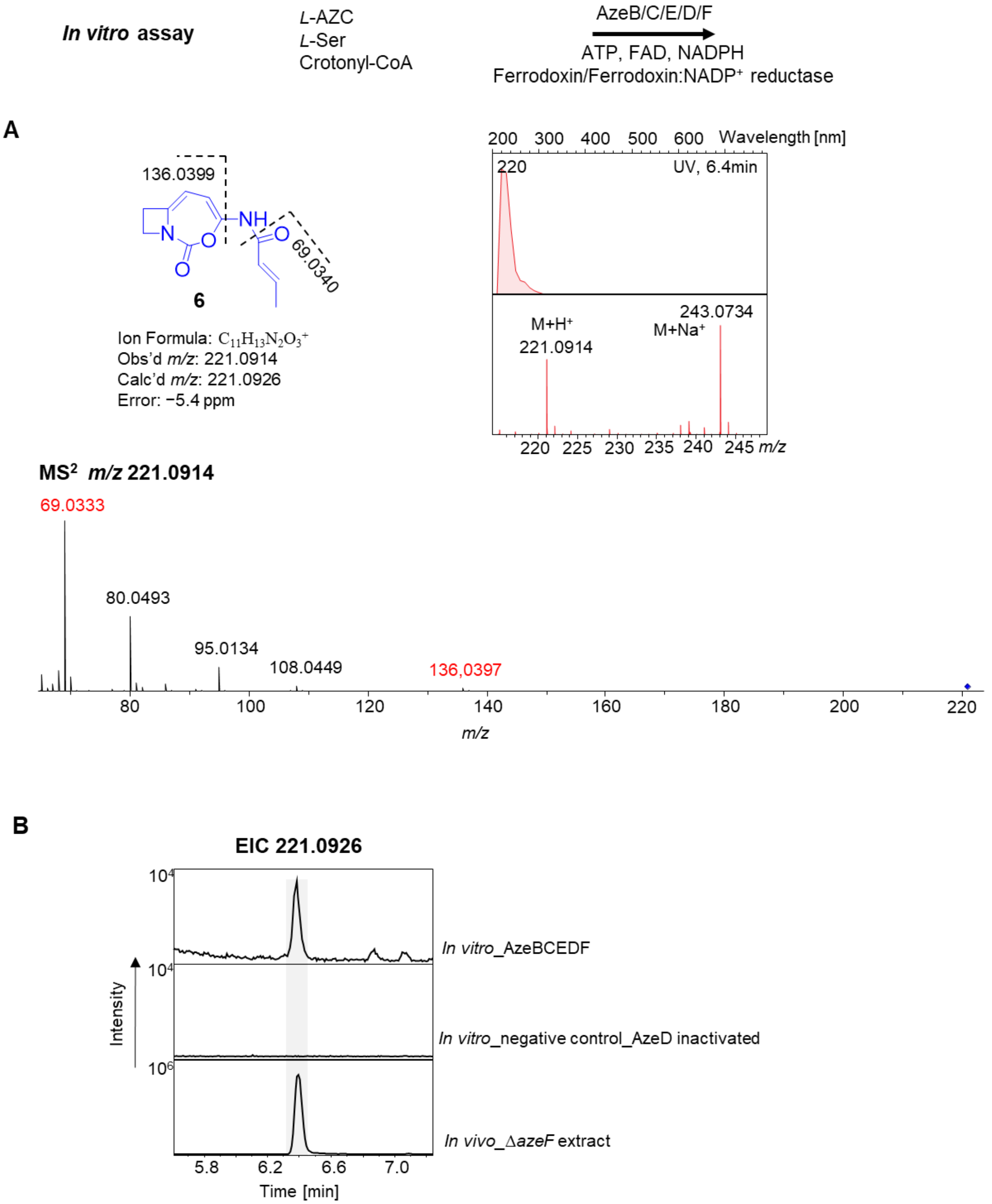


**Figure S14.** Characterization of product **6** in one-pot reaction *in vitro*. **A**) Structure, UV, MS, MS$^2$ spectra and fragment annotation of **6**. **B**) Comparison of EICs corresponding to $[M+H]^+$ ion of **6** produced *in vivo* and *in vitro*.

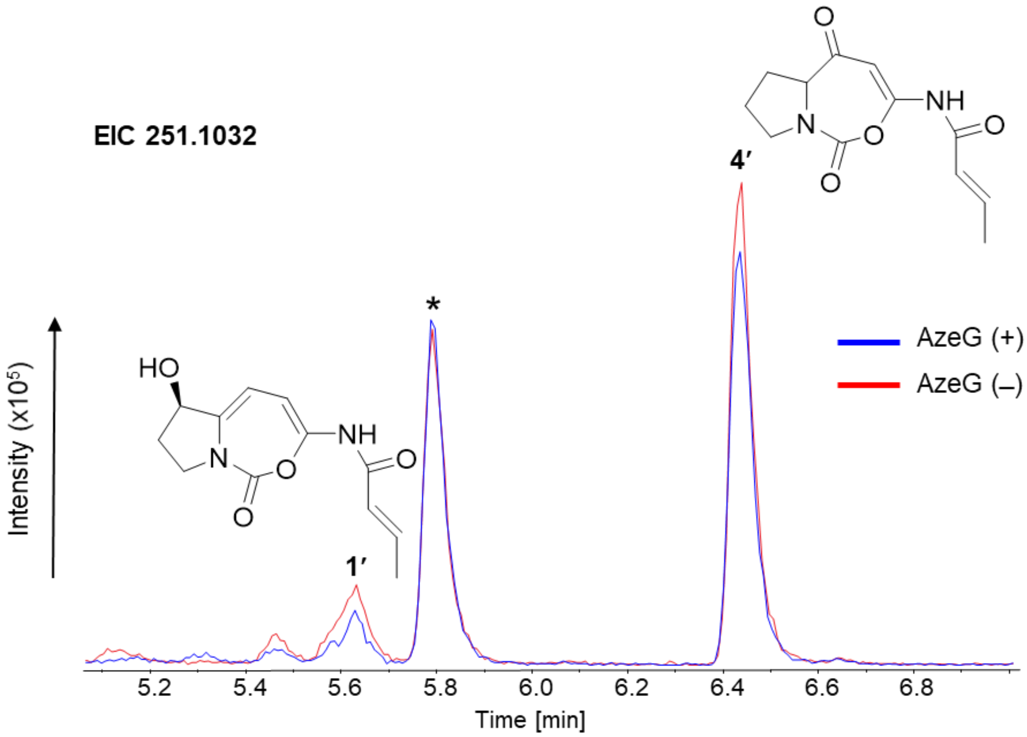


**Figure S15.** One-pot AzeB/C/E/D/F reaction *in vitro* with and without AzeG using Pro as substrate. EICs of 251.1032 corresponding to $[M+H]^+$ ion of **1′** and **4′** are shown.

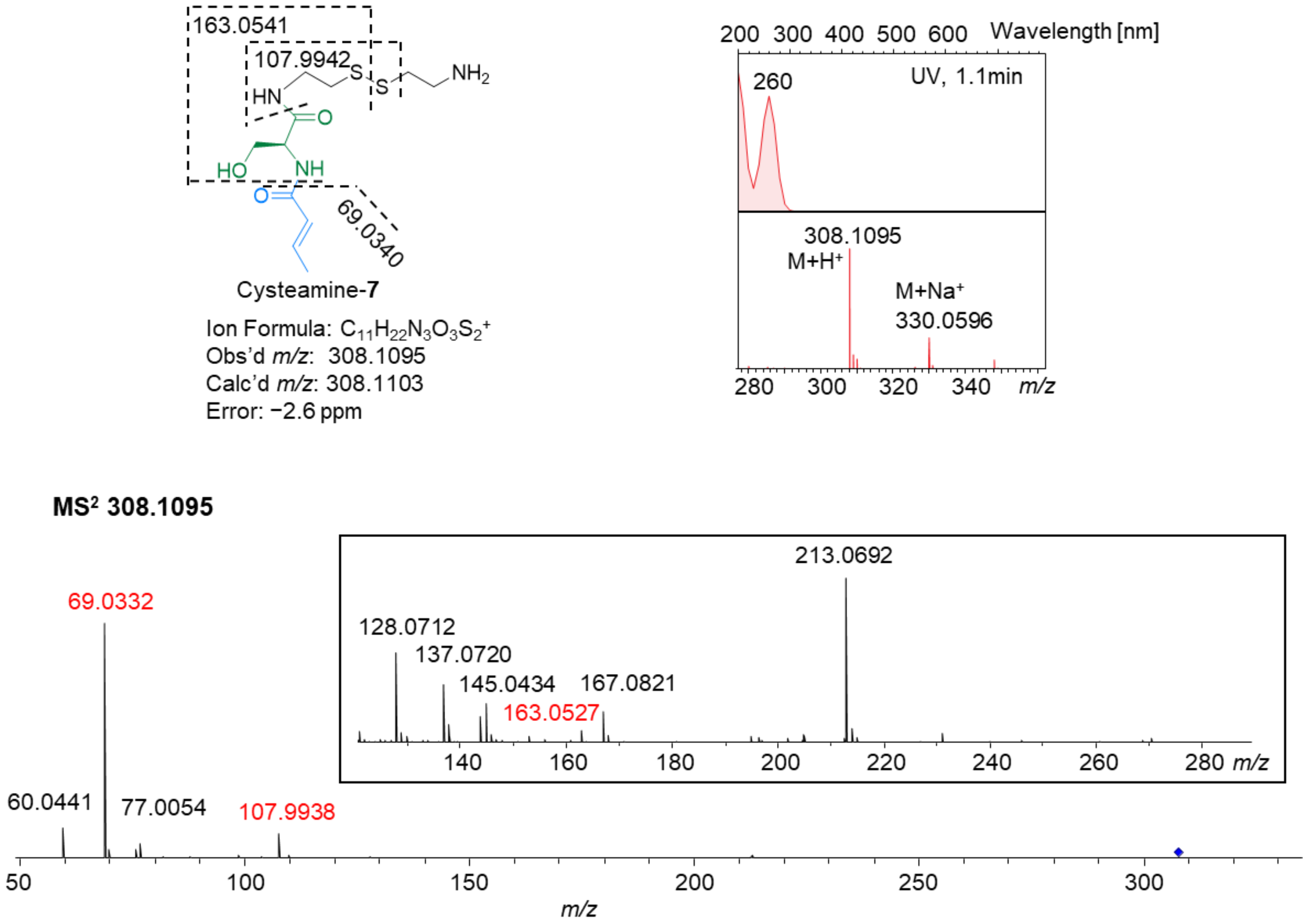


**Figure S16.** Characterization of cysteamine off-loaded intermediate **7**. The structure, UV, MS, $MS^2$ spectra and fragment annotation are shown.

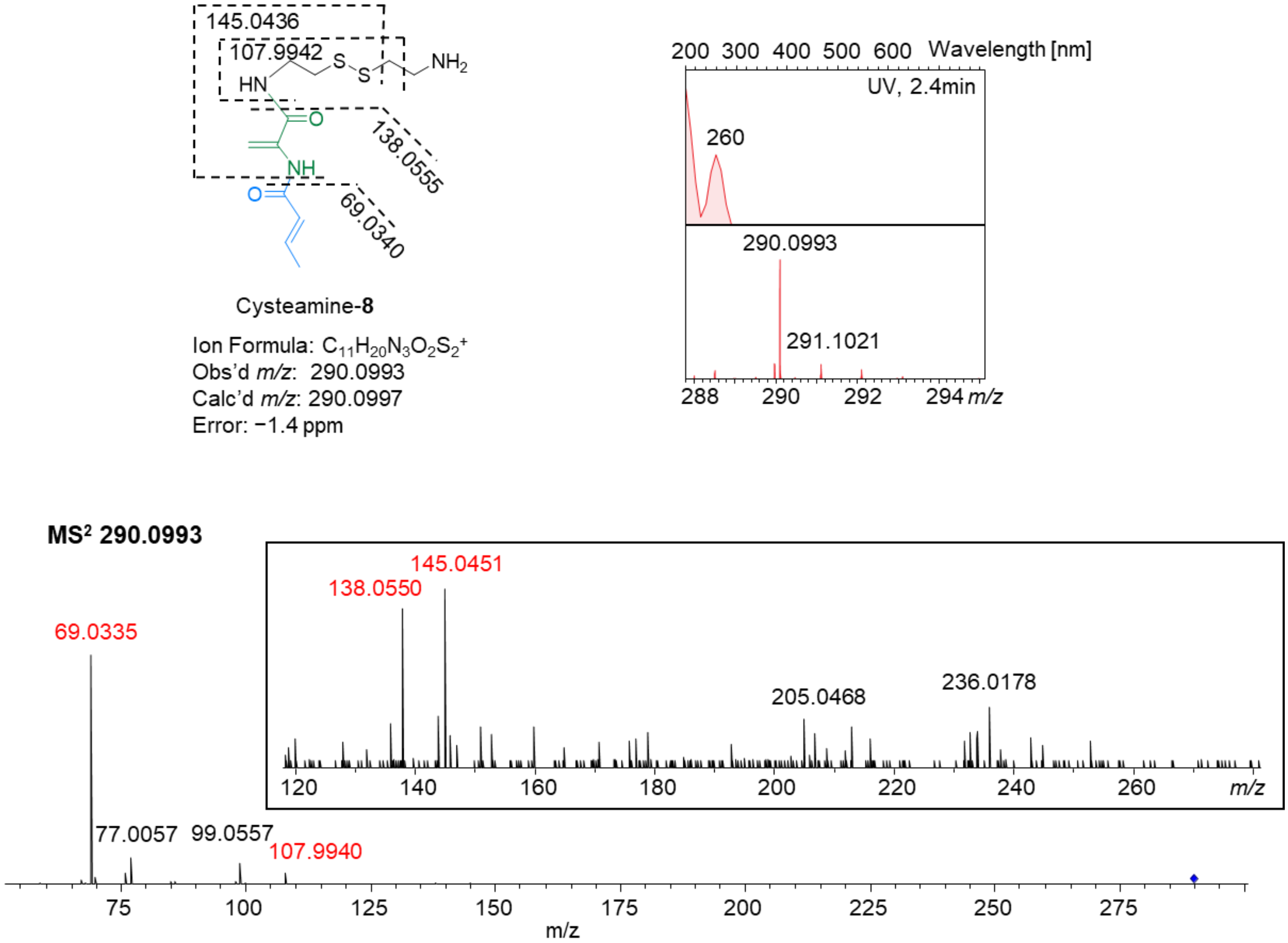


**Figure S17.** Characterization of cysteamine off-loaded intermediate **8**. The structure, UV, MS, $MS^2$ spectra and fragment annotation are shown.

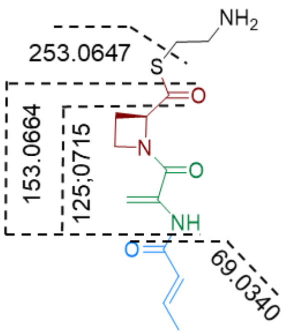


Cysteamine-**9**

Ion Formula: $C_{13}H_{20}N_3O_3S^+$
Obs'd *m/z*: 298.1218
Calc'd *m/z*: 298.1225
Error: −2.3 ppm

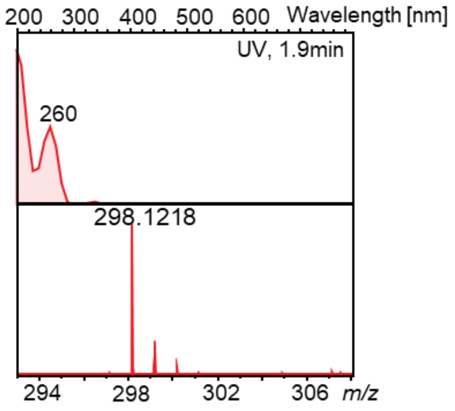


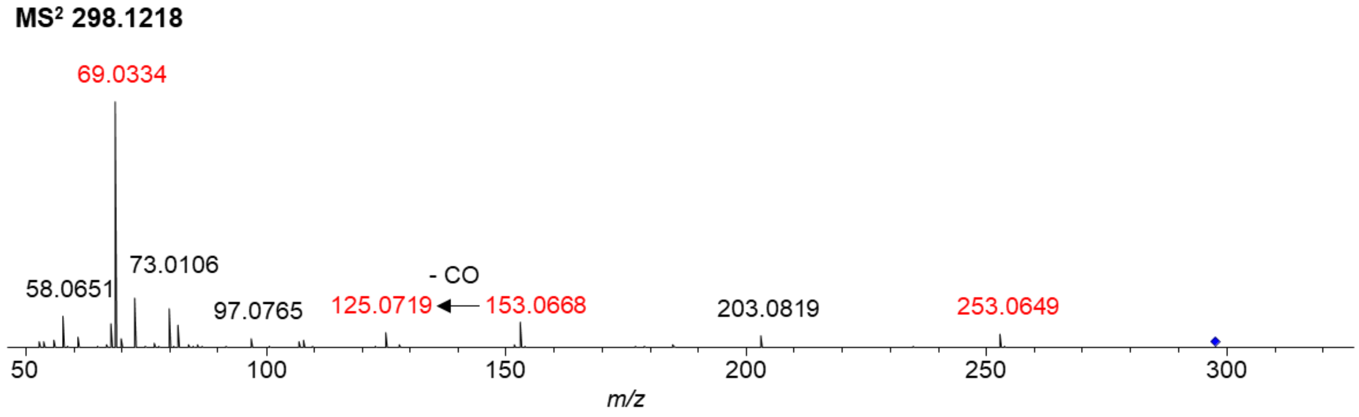


**Figure S18.** Characterization of cysteamine off-loaded intermediate **9**. The structure, UV, MS, $MS^2$ spectra and fragment annotation are shown.

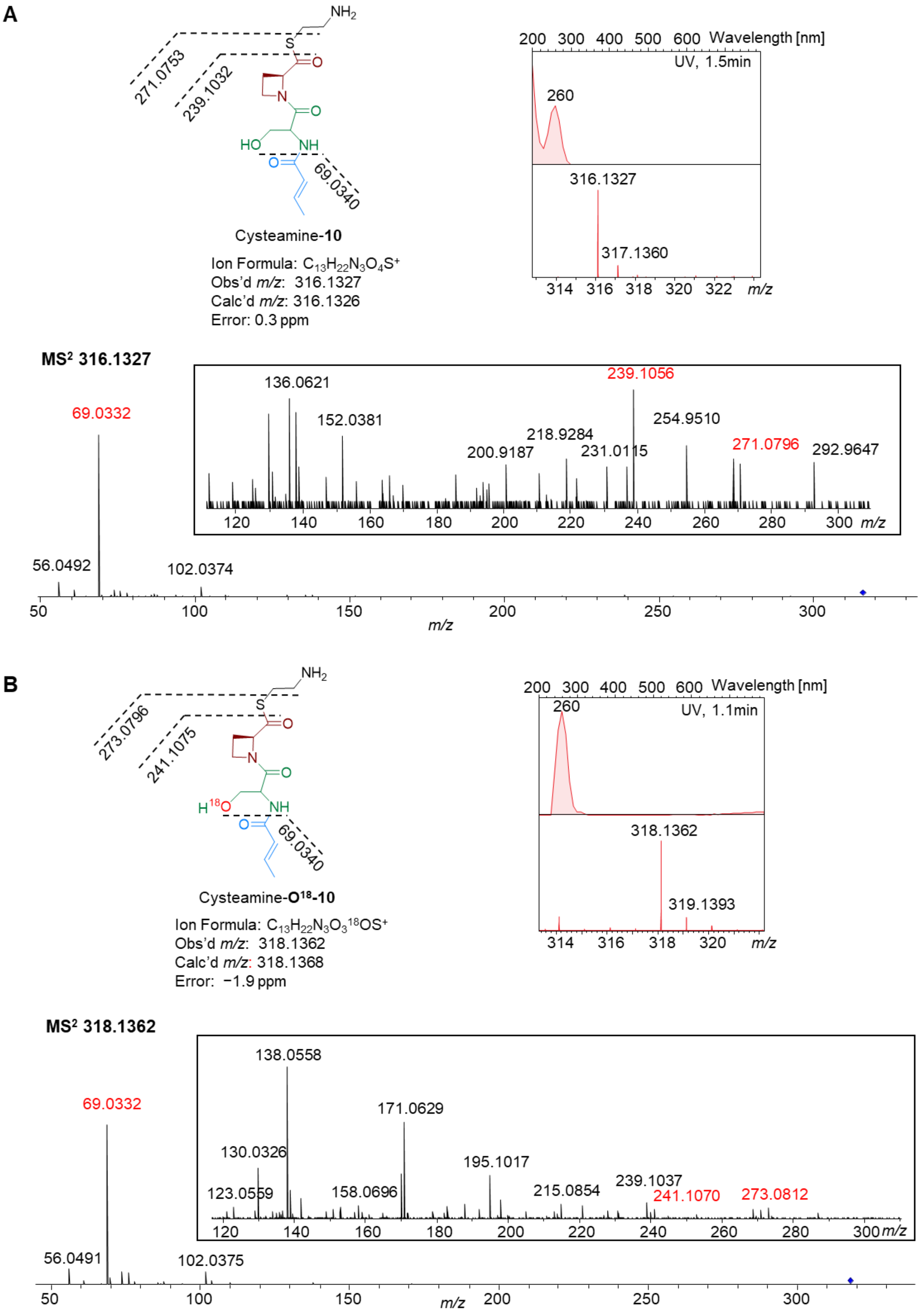


**Figure S19.** Characterization of cysteamine off-loaded intermediate **10** and its $O^{18}$-labelled product. Structures, UV, MS, $MS^2$ spectra and fragment annotation are shown.

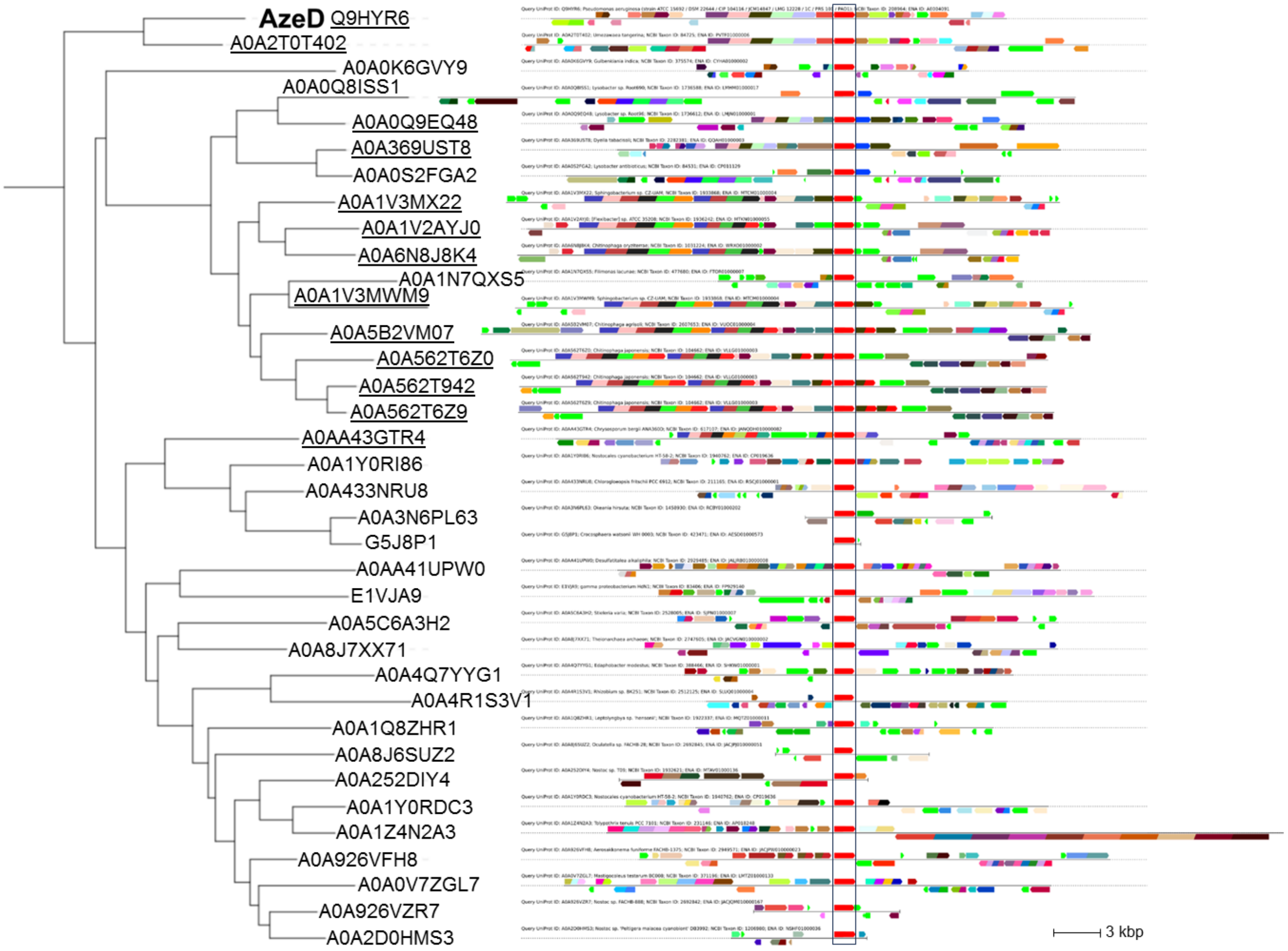


**Figure S20.** Phylogenetic tree of AzeD and homologues together with their genomic neighborhoods, as revealed by EFI-GNT analysis.[15] UniProt IDs are indicated with those surrounded by genes encoding non-ribosomal peptide synthetases or polyketide synthases underlined. In the genomic context diagram, genes encoding AzeD or homologues are boxed.

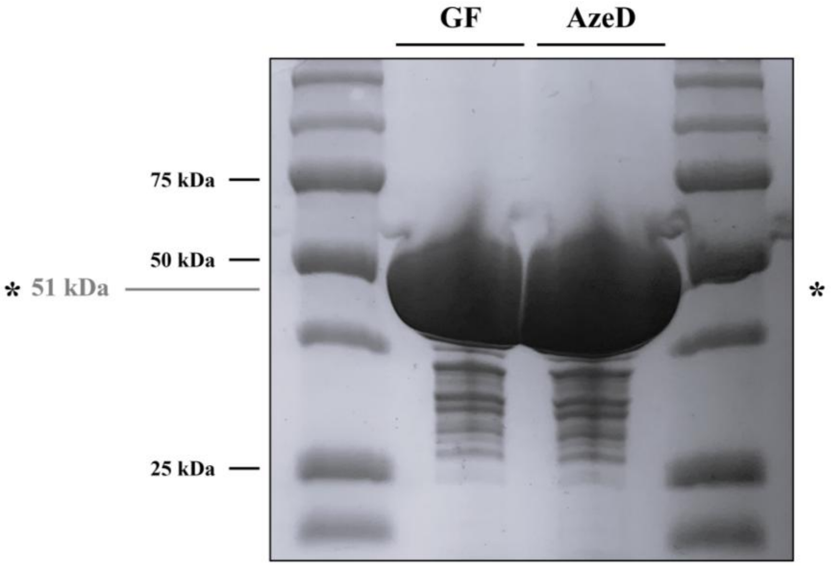


**Figure S21.** SDS-PAGE analysis of AzeD following purification. The enzyme was purified by immobilized nickel ion affinity chromatography (not shown), followed by size exclusion chromatography on a Superdex-75 column (noted GF). Due to the high calculated instability index of the protein (> 40; ProtParam tool[16,35]), the N-terminal $His_6$ tag was intentionally not cleaved prior to crystallization. The final purification yield was estimated at 34 mg/L.

### System

| | | | |
|---|---|---|---|
| Temperature (°C): | 25,0 | Duration Used (s): | 70 |
| Count Rate (kcps): | 242,5 | Measurement Position (mm): | 4,20 |
| Cell Description: | Low volume glass cuvette (1... | Attenuator: | 9 |

### Results

| | | | Size (d.n... | % Intensity: | St Dev (d.n... |
|---|---|---|---|---|---|
| Z-Average (d.nm): | 7,647 | Peak 1: | 8,260 | 100,0 | 2,323 |
| PdI: | 0,062 | Peak 2: | 0,000 | 0,0 | 0,000 |
| Intercept: | 0,934 | Peak 3: | 0,000 | 0,0 | 0,000 |
| Result quality | Good | | | | |

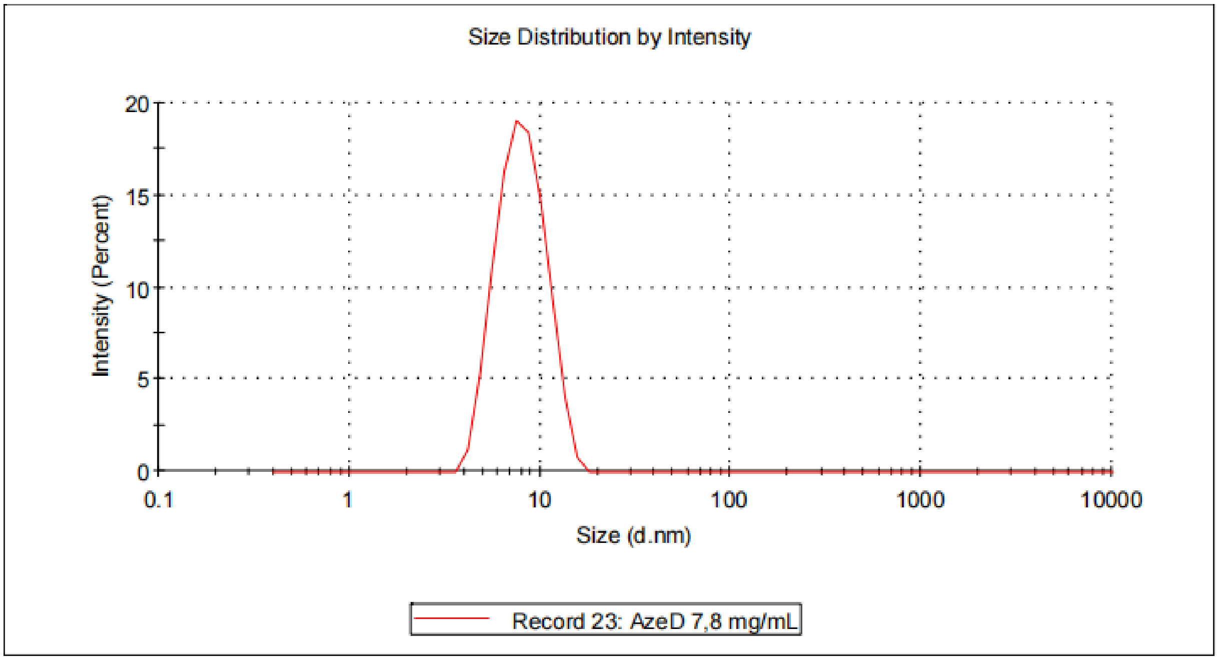


**Figure S22.** Analysis of the AzeD dehydratase by dynamic light scattering (DLS). The average hydrodynamic diameter of the proteins (noted Z-average) corresponds to 8 ± 2 nm. The sample is homogeneous with a low polydispersity index (noted PdI) of 6.2%, which is a necessary condition for further structural studies.

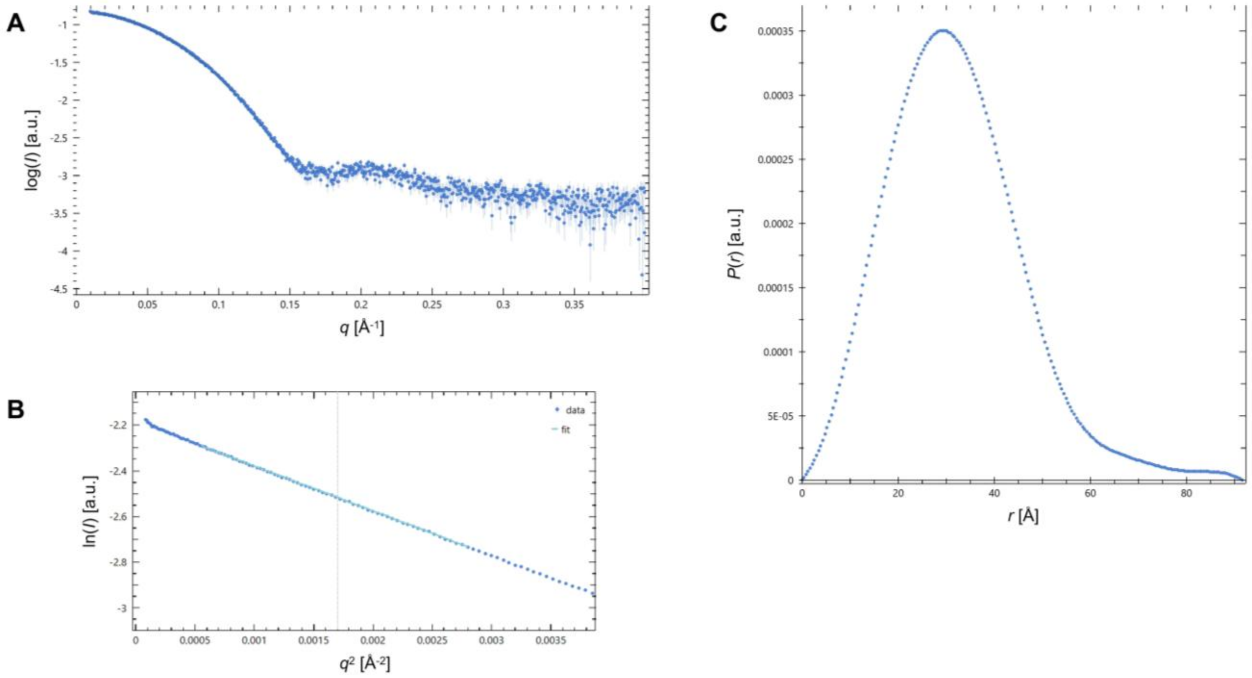


**Figure S23.** Analysis of the AzeD dehydratase by small-angle X-ray scattering (SEC-SAXS). **A**) The SAXS curve expressed as the logarithm of the diffusion intensity log($I$) of the protein as a function of the scattering vector $q$. **B**) Guinier plot which affords an initial estimate of the intensity at zero angle ($I$(0)) and the radius of gyration $R_g$. **C**) Distance distribution function ($P$($r$)), which reveals the globular shape of the protein in solution, as well as the $D_{max}$ (maximum distance) (see **Table S6**).

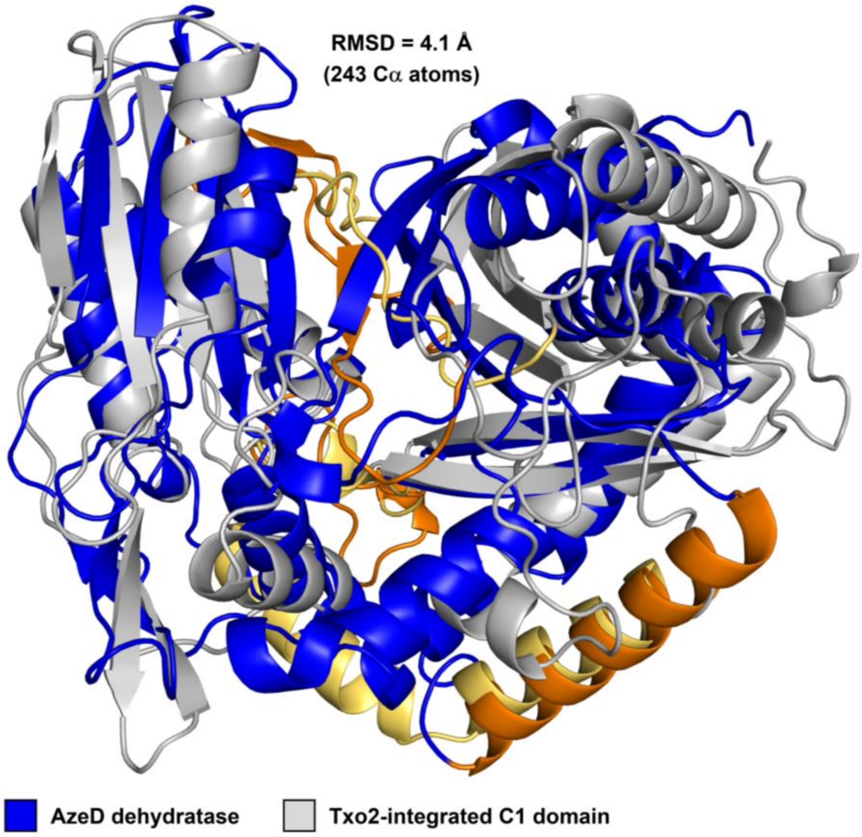
RMSD = 4.1 Å
(243 Cα atoms)
AzeD dehydratase
Txo2-integrated C1 domain

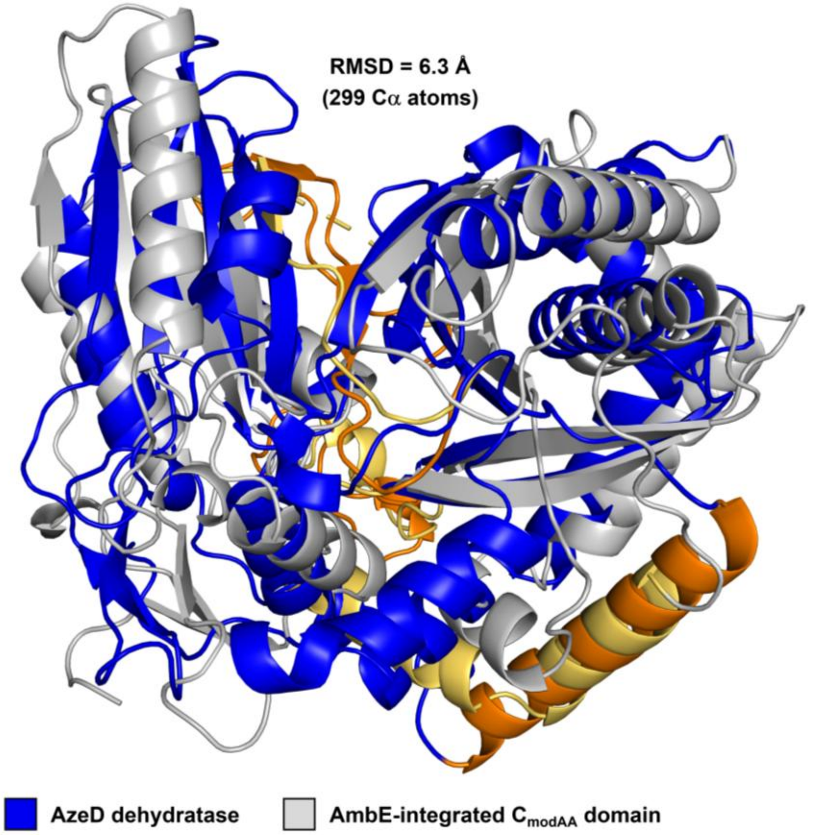
RMSD = 6.3 Å
(299 Cα atoms)
AzeD dehydratase
AmbE-integrated $C_{modAA}$ domain

**Figure S24.** Structural superimposition of AzeD, Txo2_C1 and AmbE_$C_{modAA}$ proteins. The homologues superimpose with a calculated Cα-RMSD value of 4.1 Å (243 atoms) and 6.3 Å (299 atoms), respectively, with major deviations in the degree of opening and the bridging structural elements at the interface between the N-terminal and C-terminal lobes. The AzeD dehydratase and the C domains are depicted in blue and grey, respectively, with the helix α5 and α-/β-crossovers indicated in orange (AzeD) or yellow (Txo2_C1 and AmbE_$C_{modAA}$).

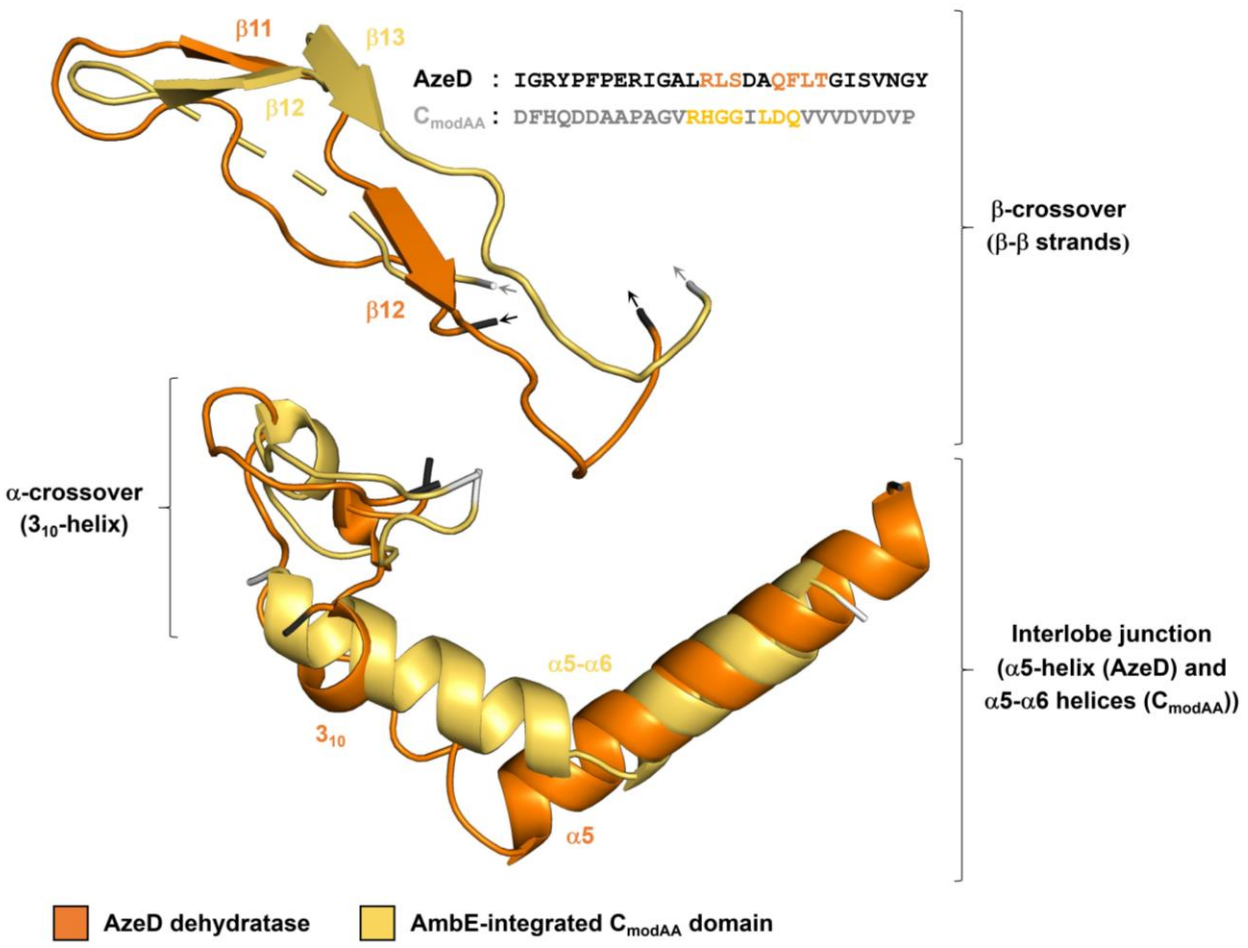


**Figure S25.** Comparison between AzeD and AmbE_$C_{modAA}$ of the organization of the bridging structural elements at the interlobe interface. The bridging α-helices and α-/β-crossovers of the AzeD dehydratase and AmbE_$C_{modAA}$ domain, represented in orange and yellow, respectively, form the junction between the N- and C-terminal lobes within both structures. Notable differences include the linker region, as a residual $3_{10}$-helix is present in AzeD instead of a classical α-helix, and the relative length of the adjacent helix (α5 (AzeD) and α6 (AmbE_$C_{modAA}$)), as four additional turns are present in AzeD α5. Further divergence is evident in the orientation of helix $3_{10}$ of the α-crossover, as well as in the length and residue composition of the extended β-crossover (i.e. amino acids (QDDAA) are absent in the homologue).

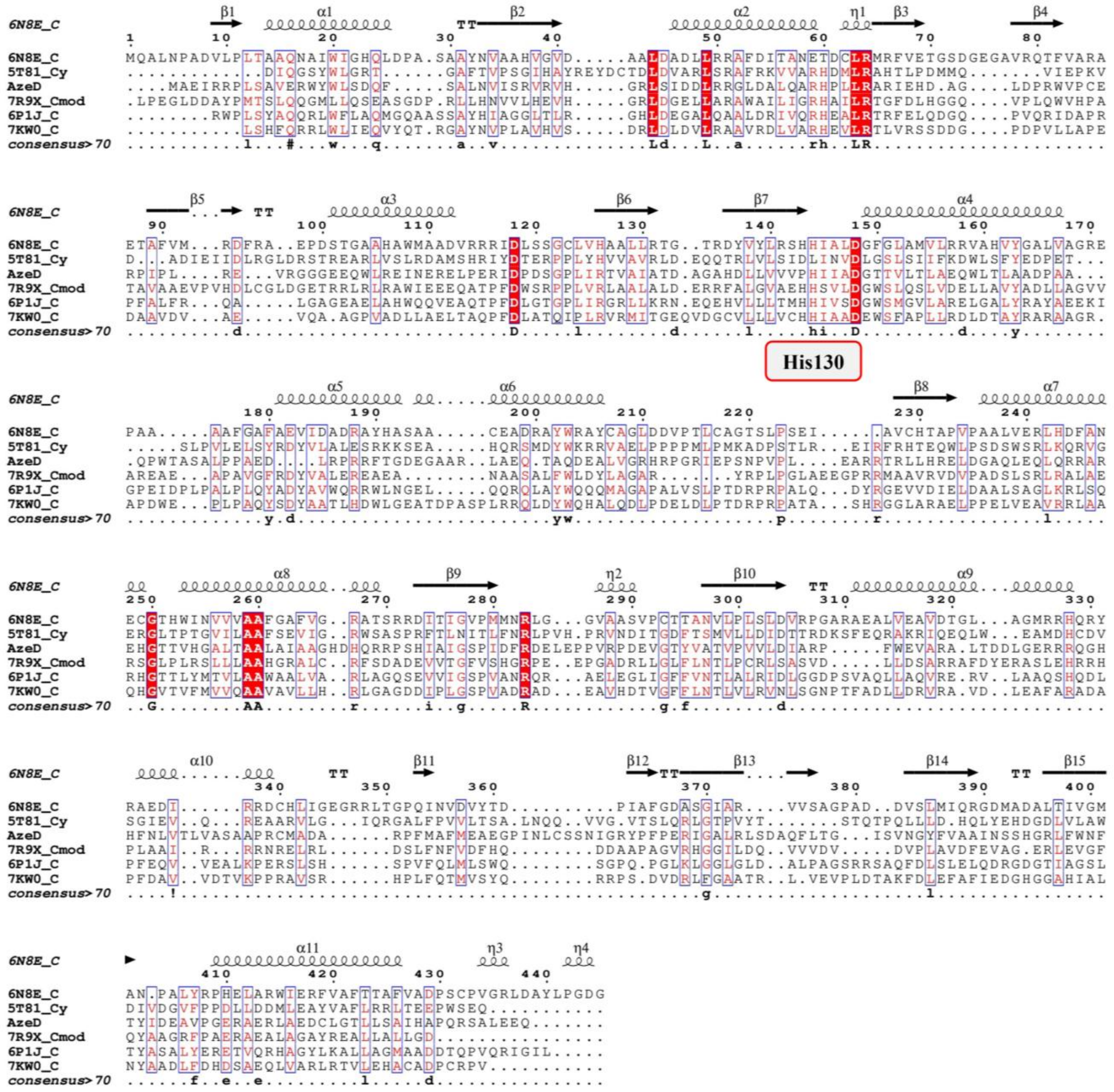

**Figure S26.** Sequence alignment of AzeD dehydratase homologues. The protein sequence of AzeD was aligned against sequences of NRPS-integrated C domains with condensation (PDB ID: 6P1J, 7KW0 and 6N8E), heterocyclization (PDB: 5T81) or atypical dehydration functions (PDB ID: 7R9X). NRPS C domains were identified using the PDB database[36] and the Dali structural comparison server.[37] The AzeD dehydratase and the related C domains exhibit a sequence identity ranging from 17%–22% and share determinants involved in substrate binding and the catalytic reaction, including residues His130 and Asp134 of the consensus motif xHxxxDx.

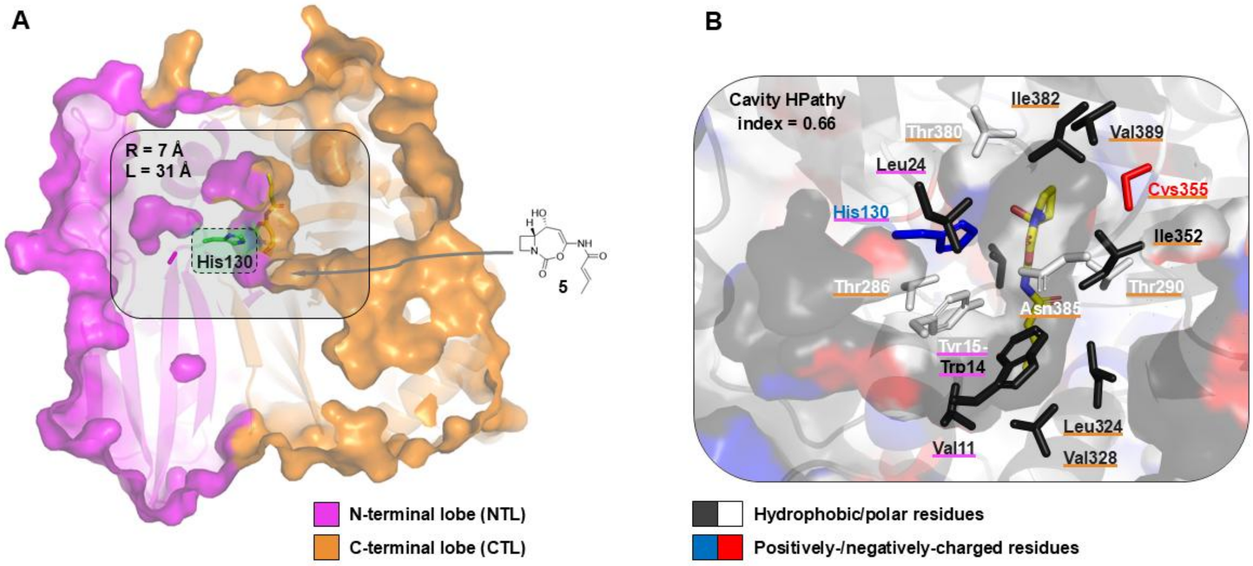


**Figure S27.** Shape and amino acid composition of the active site of the AzeD dehydratase in the presence of computationally docked **5**. **A**) The N-terminal and C-terminal domains, depicted in magenta and orange, respectively, create an extended active site at their interface. The lone entry tunnel has dimensions R = 7 Å and L = 31 Å. The catalytic His130 is buried at a distance of approximately 11 Å from the protein surface. **B**) The active site cavity is largely hydrophobic with a positive hydropathy index (0.66) (calculated using the PDBsum program[38]). Multiple aliphatic, aromatic and charged residues furnished by both lobes contribute to ligand binding. Hydrophobic, polar, positively- and negatively-charged residues are indicated in black, white, blue, and red, respectively.

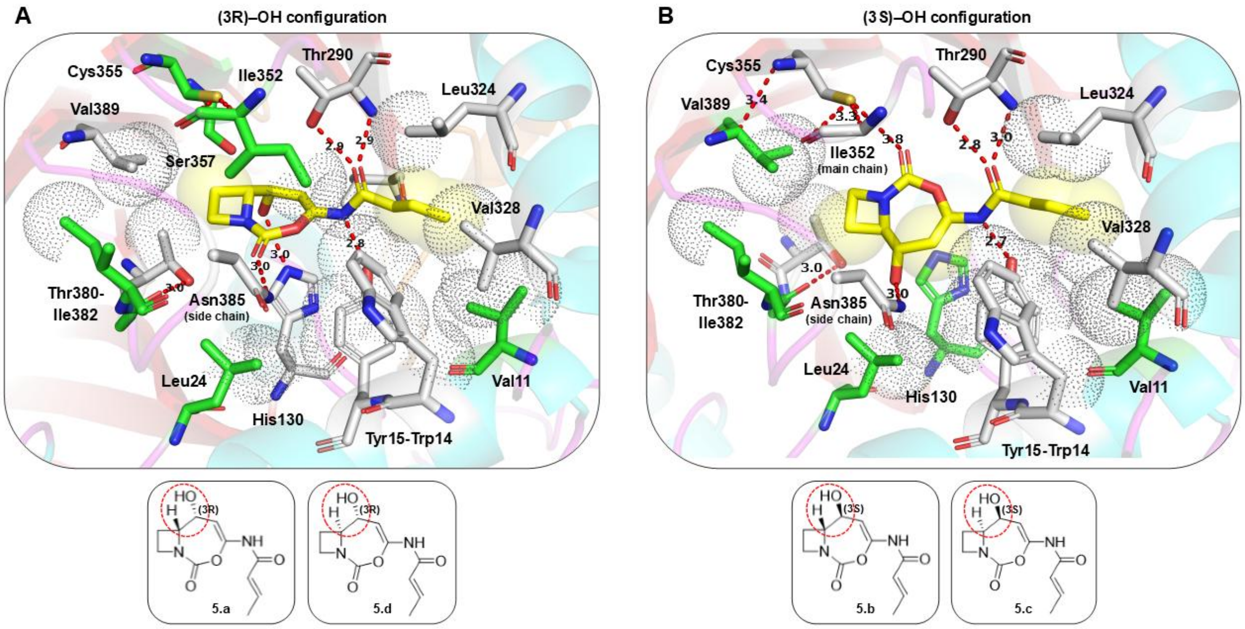


**Figure S28.** Molecular docking analysis of four stereoisomers of **5** in the active site of AzeD. The same docking pose is observed for **5.a** (shown) and **5.d** for which the 3-OH is in the *R*-configuration, while **5.b** (shown) and **5.c** which both incorporate a (3*S*)-OH, also adopt the same pose.

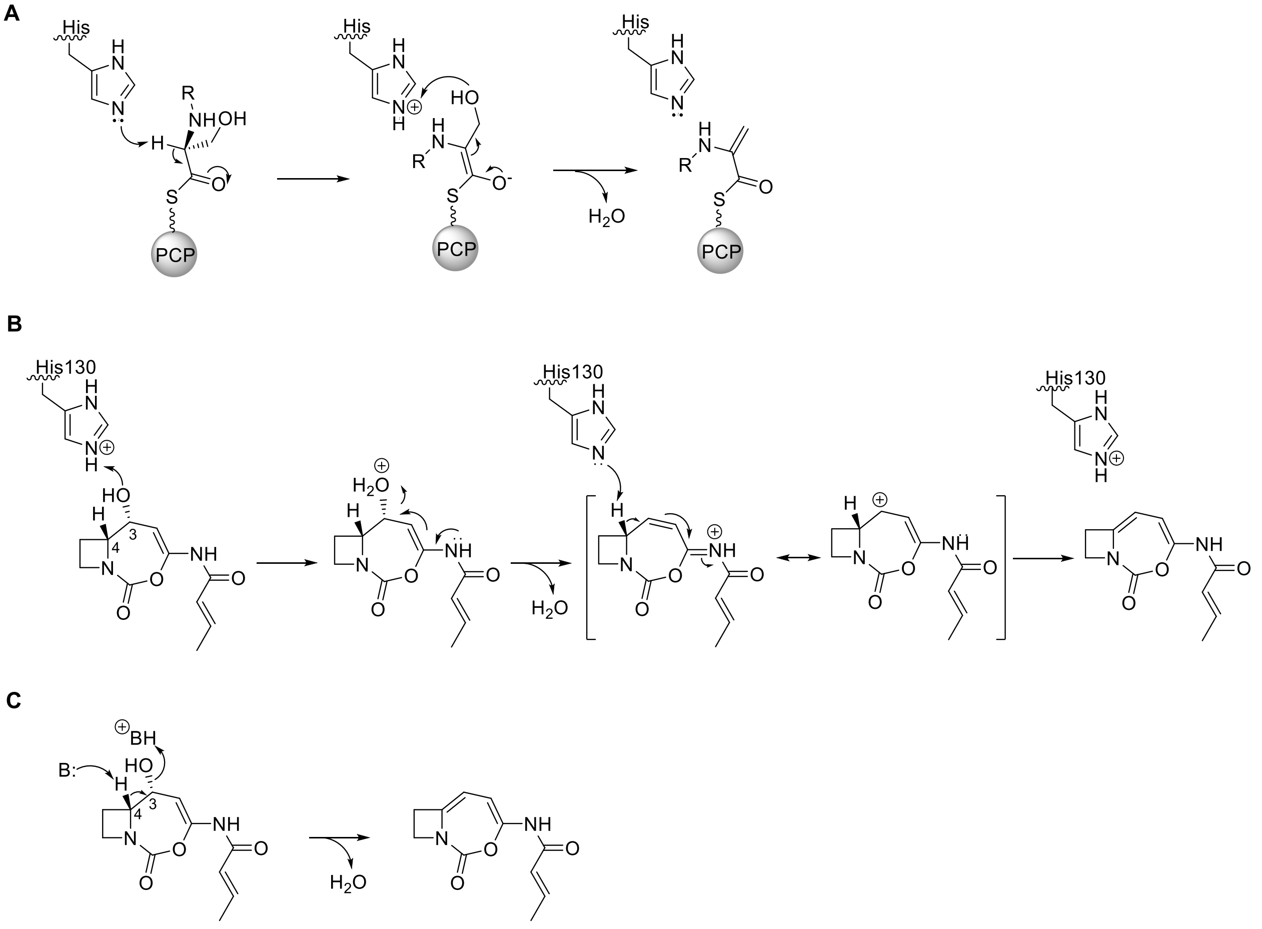


**Figure S29**. Possible mechanisms for C domains or proteins-catalyzed dehydration. **A**) Proposed E1cB mechanism for online dehydration performed by NRPS C domains[39,40]. The active site His functions as a general base for α-proton abstraction to yield an enolate, then as a general acid for dehydration. **B**) Proposed E1 mechanism for AzeD-mediated dehydration based on both crystallographic and mutagenesis data. In this mechanism, His130 that was shown to be critical by site-directed mutagenesis, first acts as a general acid to activate the water leaving group, and then as a general base to initiate the double bond migration to give the final product. **C**) Alternative E2 mechanism for AzeD reaction. In this proposal, the dehydration occurs in concerted fashion catalyzed by the synchronous action of a general acid and base. The *anti*-periplanar stereochemistry required for this reaction would occur naturally if the hydroxyl leaving group at C-3 exhibits R stereochemistry (the R stereochemistry of the C4-H was established previously in azetidomonamide B **2**[3]).

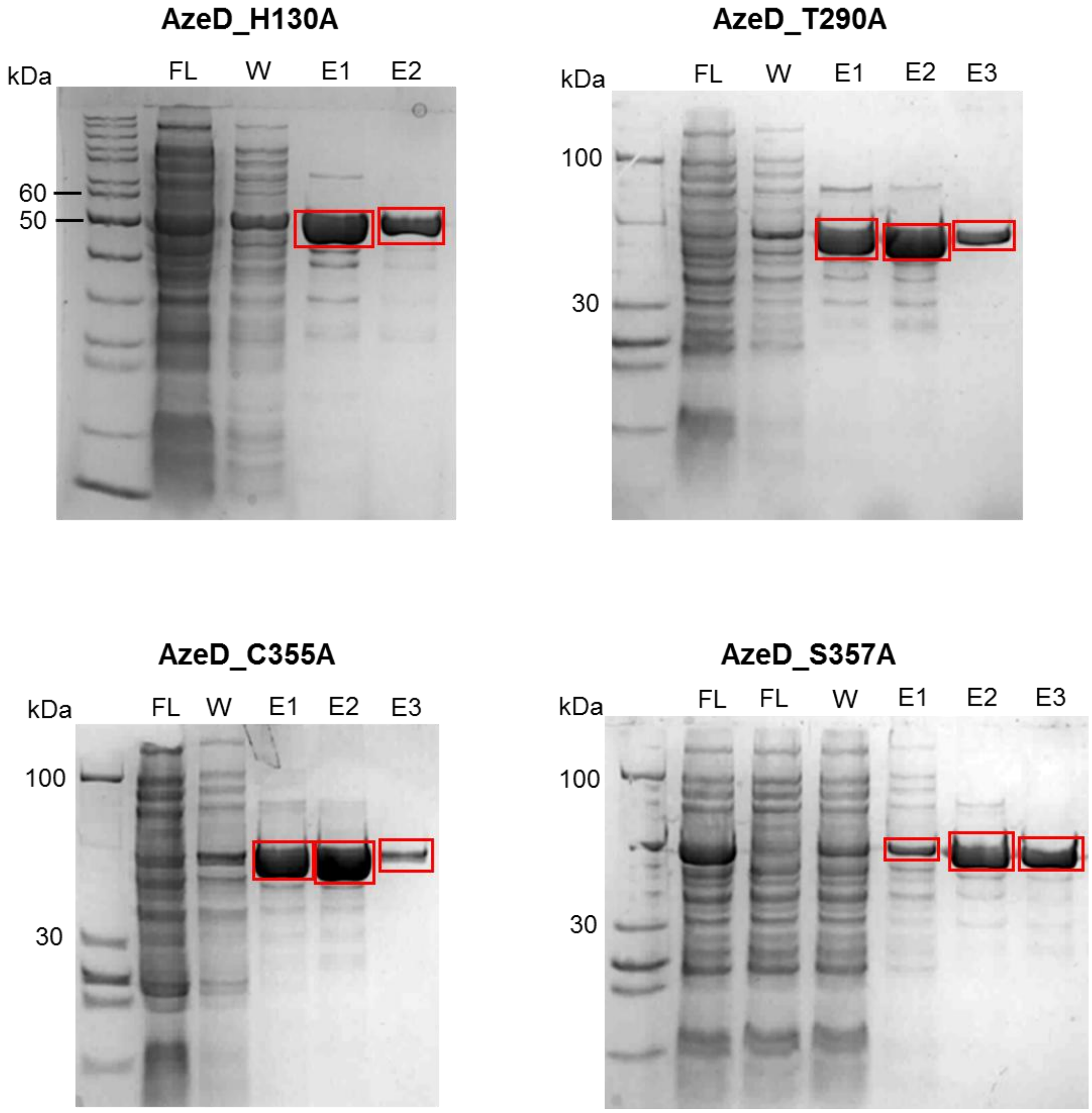


**Figure S30.** SDS-PAGE analysis of AzeD mutant purification by nickel affinity chromatography. FL, flowthrough; W, wash; E, elution. The target proteins in the elution fractions are boxed.

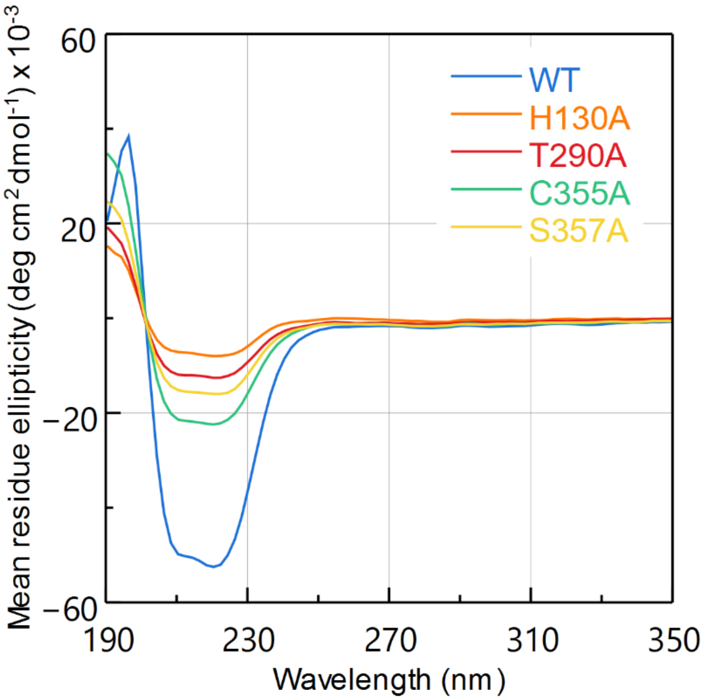


**Figure S31.** Circular dichroism spectra of AzeD wild-type and mutant proteins. Mean residue ellipticity is plotted as a function of wavelength. These data indicate varying degree of structural destabilization induced by mutations, with H130A having the most substantial impact.

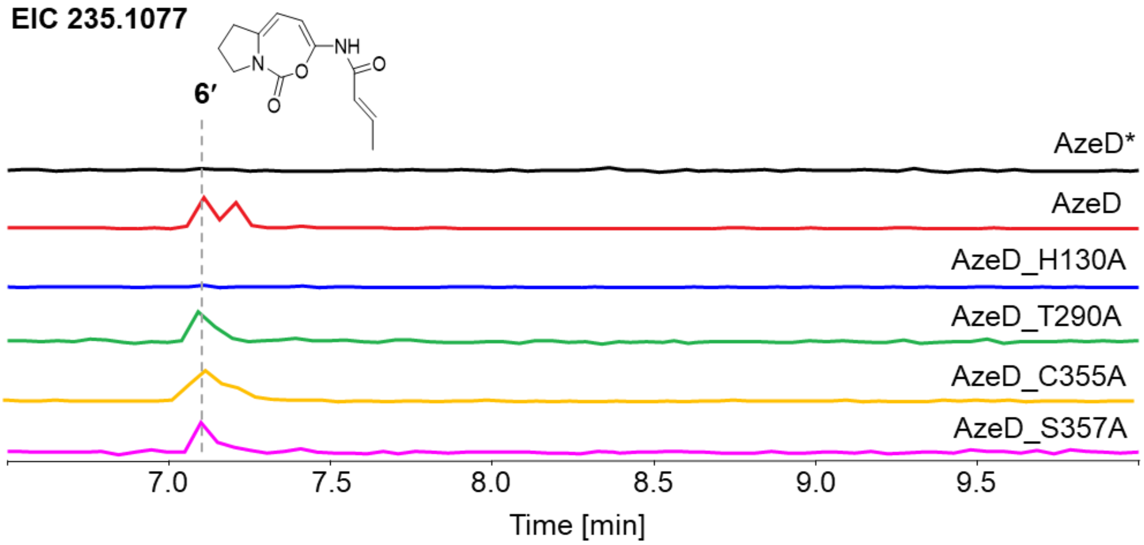


**Figure S32.** Activities of AzeD mutants in one-pot AzeB/C/E/D reaction using Pro as substrate. EICs of 235.1077 corresponding to $[M+H]^+$ ion of **6′** are shown. Substitution of H130 to A completely abolished the dehydration activity. * denotes heat-inactivated enzyme.

(13) Guindon, S.; Dufayard, J.-F.; Lefort, V.; Anisimova, M.; Hordijk, W.; Gascuel, O. New

Algorithms and Methods to Estimate Maximum-Likelihood Phylogenies: Assessing the Performance of PhyML 3.0. *Syst. Biol.* **2010**, *59* (3), 307–321. https://doi.org/10.1093/sysbio/syq010.

(21) Jumper, J.; Evans, R.; Pritzel, A.; Green, T.; Figurnov, M.; Ronneberger, O.; Tunyasuvunakool, K.; Bates, R.; Žídek, A.; Potapenko, A.; Bridgland, A.; Meyer, C.; Kohl, S. A. A.; Ballard, A. J.; Cowie, A.; Romera-Paredes, B.; Nikolov, S.; Jain, R.; Adler, J.; Back, T.; Petersen, S.; Reiman, D.; Clancy, E.; Zielinski, M.; Steinegger, M.; Pacholska, M.; Berghammer, T.; Bodenstein, S.; Silver, D.; Vinyals, O.; Senior, A. W.; Kavukcuoglu, K.; Kohli, P.; Hassabis, D. Highly Accurate Protein Structure Prediction with AlphaFold. *Nature* **2021**, *596* (7873), 583–589. https://doi.org/10.1038/s41586-021-03819-2.

(22) Emsley, P.; Lohkamp, B.; Scott, W. G.; Cowtan, K. Features and Development of *Coot*. *Acta Crystallogr. D Biol. Crystallogr.* **2010**, *66* (4), 486–501. https://doi.org/10.1107/S0907444910007493.

(23) Murshudov, G. N.; Skubák, P.; Lebedev, A. A.; Pannu, N. S.; Steiner, R. A.; Nicholls, R. A.; Winn, M. D.; Long, F.; Vagin, A. A. *REFMAC* 5 for the Refinement of Macromolecular Crystal Structures. *Acta Crystallogr. D Biol. Crystallogr.* **2011**, *67* (4), 355–367. https://doi.org/10.1107/S0907444911001314.

(24) Chen, V. B.; Arendall, W. B.; Headd, J. J.; Keedy, D. A.; Immormino, R. M.; Kapral, G. J.; Murray, L. W.; Richardson, J. S.; Richardson, D. C. *MolProbity* : All-Atom Structure Validation for Macromolecular Crystallography. *Acta Crystallogr. D Biol. Crystallogr.* **2010**,

*66* (1), 12–21. https://doi.org/10.1107/S0907444909042073.
(25) Young, J. Y.; Westbrook, J. D.; Feng, Z.; Sala, R.; Peisach, E.; Oldfield, T. J.; Sen, S.; Gutmanas, A.; Armstrong, D. R.; Berrisford, J. M.; Chen, L.; Chen, M.; Di Costanzo, L.; Dimitropoulos, D.; Gao, G.; Ghosh, S.; Gore, S.; Guranovic, V.; Hendrickx, P. M. S.; Hudson, B. P.; Igarashi, R.; Ikegawa, Y.; Kobayashi, N.; Lawson, C. L.; Liang, Y.; Mading, S.; Mak, L.; Mir, M. S.; Mukhopadhyay, A.; Patwardhan, A.; Persikova, I.; Rinaldi, L.; Sanz-Garcia, E.; Sekharan, M. R.; Shao, C.; Swaminathan, G. J.; Tan, L.; Ulrich, E. L.; Van Ginkel, G.; Yamashita, R.; Yang, H.; Zhuravleva, M. A.; Quesada, M.; Kleywegt, G. J.; Berman, H. M.; Markley, J. L.; Nakamura, H.; Velankar, S.; Burley, S. K. OneDep: Unified wwPDB System for Deposition, Biocuration, and Validation of Macromolecular Structures in the PDB Archive. *Structure* **2017**, *25* (3), 536–545. https://doi.org/10.1016/j.str.2017.01.004.
(26) Liu, Y.; Yang, X.; Gan, J.; Chen, S.; Xiao, Z.-X.; Cao, Y. CB-Dock2: Improved Protein–Ligand Blind Docking by Integrating Cavity Detection, Docking and Homologous Template Fitting. *Nucleic Acids Res.* **2022**, *50* (W1), W159–W164. https://doi.org/10.1093/nar/gkac394.
(27) Lilkova, E.; Petkov, P.; Ilieva, N.; Litov, L. The PyMOL Molecular Graphics System, Version 2.0 Schrodinger, LLC. 2015, 10919-10925., 2015.
(28) Thureau, A.; Roblin, P.; Pérez, J. BioSAXS on the SWING Beamline at Synchrotron SOLEIL. *J. Appl. Crystallogr.* **2021**, *54* (6), 1698–1710. https://doi.org/10.1107/s1600576721008736.
(29) Franke, D.; Petoukhov, M. V.; Konarev, P. V.; Panjkovich, A.; Tuukkanen, A.; Mertens, H. D. T.; Kikhney, A. G.; Hajizadeh, N. R.; Franklin, J. M.; Jeffries, C. M.; Svergun, D. I. *ATSAS 2.8*: A Comprehensive Data Analysis Suite for Small-Angle Scattering from Macromolecular Solutions. *J. Appl. Crystallogr.* **2017**, *50* (4), 1212–1225. https://doi.org/10.1107/s1600576717007786.
(30) Svergun, D. I. Determination of the Regularization Parameter in Indirect-Transform Methods Using Perceptual Criteria. *J. Appl. Crystallogr.* **1992**, *25* (4), 495–503. https://doi.org/10.1107/s0021889892001663.
(31) Manalastas-Cantos, K.; Konarev, P. V.; Hajizadeh, N. R.; Kikhney, A. G.; Petoukhov, M. V.; Molodenskiy, D. S.; Panjkovich, A.; Mertens, H. D. T.; Gruzinov, A.; Borges, C.; Jeffries, C. M.; Svergun, D. I.; Franke, D. *ATSAS 3.0*: Expanded Functionality and New Tools for Small-Angle Scattering Data Analysis. *J. Appl. Crystallogr.* **2021**, *54* (1), 343–355. https://doi.org/10.1107/s1600576720013412.
(32) Liu, Y.; Yang, X.; Gan, J.; Chen, S.; Xiao, Z.-X.; Cao, Y. CB-Dock2: Improved Protein–Ligand Blind Docking by Integrating Cavity Detection, Docking and Homologous Template Fitting. *Nucleic Acids Res.* **2022**, *50* (W1), W159–W164. https://doi.org/10.1093/nar/gkac394.
(33) Herrero, M.; De Lorenzo, V.; Timmis, K. N. Transposon Vectors Containing Non-Antibiotic Resistance Selection Markers for Cloning and Stable Chromosomal Insertion of Foreign Genes in Gram-Negative Bacteria. *J. Bacteriol.* **1990**, *172* (11), 6557–6567. https://doi.org/10.1128/jb.172.11.6557-6567.1990.
(34) Lacks, S.; Greenberg, B. Complementary Specificity of Restriction Endonucleases of Diplococcus Pneumoniae with Respect to DNA Methylation. *J. Mol. Biol.* **1977**, *114* (1), 153–168. https://doi.org/10.1016/0022-2836(77)90289-3.
(35) Gamage, D. G.; Gunaratne, A.; Periyannan, G. R.; Russell, T. G. Applicability of Instability

Index for In Vitro Protein Stability Prediction. *Protein Pept. Lett.* **2019**, *26* (5), 339–347. https://doi.org/10.2174/0929866526666190228144219.
